# Impact of higher-order network structure on emergent cortical activity

**DOI:** 10.1101/802074

**Authors:** Max Nolte, Eyal Gal, Henry Markram, Michael W. Reimann

## Abstract

Synaptic connectivity between neocortical neurons is highly structured. The network structure of synaptic connectivity includes first-order properties that can be described by pairwise statistics, such as strengths of connections between different neuron types and distance-dependent connectivity, and higher-order properties, such as an abundance of cliques of all-to-all connected neurons. The relative impact of first- and higher-order structure on emergent cortical network activity is unknown. Here, we compare network structure and emergent activity in two neocortical microcircuit models with different synaptic connectivity. Both models have a similar first-order structure, but only one model includes higher-order structure arising from morphological diversity within neuronal types. We find that such morphological diversity leads to more heterogeneous degree distributions, increases the number of cliques, and contributes to a small-world topology. The increase in higher-order network structure is accompanied by more nuanced changes in neuronal firing patterns, such as an increased dependence of pairwise correlations on the positions of neurons in cliques. Our study shows that circuit models with very similar first-order structure of synaptic connectivity can have a drastically different higher-order network structure, and suggests that the higher-order structure imposed by morphological diversity within neuronal types has an impact on emergent cortical activity.

## INTRODUCTION

Local synaptic connectivity between neocortical neurons is highly structured (Perin, Berger, & Markram, 2011; Song, Sjstrm, Reigl, Nelson, & Chklovskii, 2005). Details of *first-order structure* that can be described by pairwise statistics include distinct mean connection strengths between different neuron types (Feldmeyer, Lubke, Silver, & Sakmann, 2002; Jiang et al., 2015; Le B, Silberberg, Wang, & Markram, 2007; Silberberg & Markram, 2007), distance-dependent connectivity that changes between neuron types (Fino & Yuste, 2011; Holmgren, Harkany, Svennenfors, & Zilberter, 2003; Jiang et al., 2015; Song et al., 2005), and a bias for reciprocal connections (Markram et al., 2015; Perin et al., 2011; Song et al., 2005). This first-order structure is undoubtedly important for emergent electrical activity, for example by constraining the interlaminar flow of spiking activity (Reyes-Puerta, Sun, Kim, Kilb, & Luhmann, 2014) and constraining the excitation-inhibition balance (Rosenbaum, Smith, Kohn, Rubin, & Doiron, 2017).

Yet, local synaptic connectivity also contains significant *higher-order structure* that cannot be described by pairwise statistics (Benson, Gleich, & Leskovec, 2016). Examples are an overexpression of certain triplet motifs of neurons (Perin et al., 2011; Song et al., 2005) and an abundance of cliques of all-to-all connected neurons (Reimann, Nolte, et al., 2017). Such higher-order structure has been hypothesized to be important for computation (Braitenberg, 1978; Hebb, 1949; Knoblauch, Palm, & Sommer, 2009; Willshaw, Buneman, & Longuet-Higgins, 1969). On the other hand, modern artificial neural networks have demonstrated impressive computational capabilities without complex higher-order micro-structures (Simonyan & Zisserman, 2014). Whether computation in the cortex relies on higher-order structure such as multi-neuron motifs on top of already complex first-order structure is unknown.

Answering this question in vivo will require simultaneous access to both detailed synaptic connectivity and electrical activity. While detailed synaptic connectivity of larger areas encompassing thousands of neurons might soon become available (Kasthuri et al., 2015; Yin et al., 2019), it will remain difficult to study the direct impact of the network structure on electrical activity, and even then it would be difficult to quantify the relative impact of first- and higher-order structure. A modelling approach can help bridge this gap. An algorithmic approach uses available data to generate synaptic connectivity in a neocortical microcircuit model with diverse morphologies (Reimann, King, Muller, Ramaswamy, & Markram, 2015). When simulated, this neocortical microcircuit model (*NMC-model*, Figure 1A1) can reproduce an array of in vivo-like neuronal activity (Markram et al., 2015), and allow us to compare and manipulate detailed, predicted structure and function.

**Figure 1.**
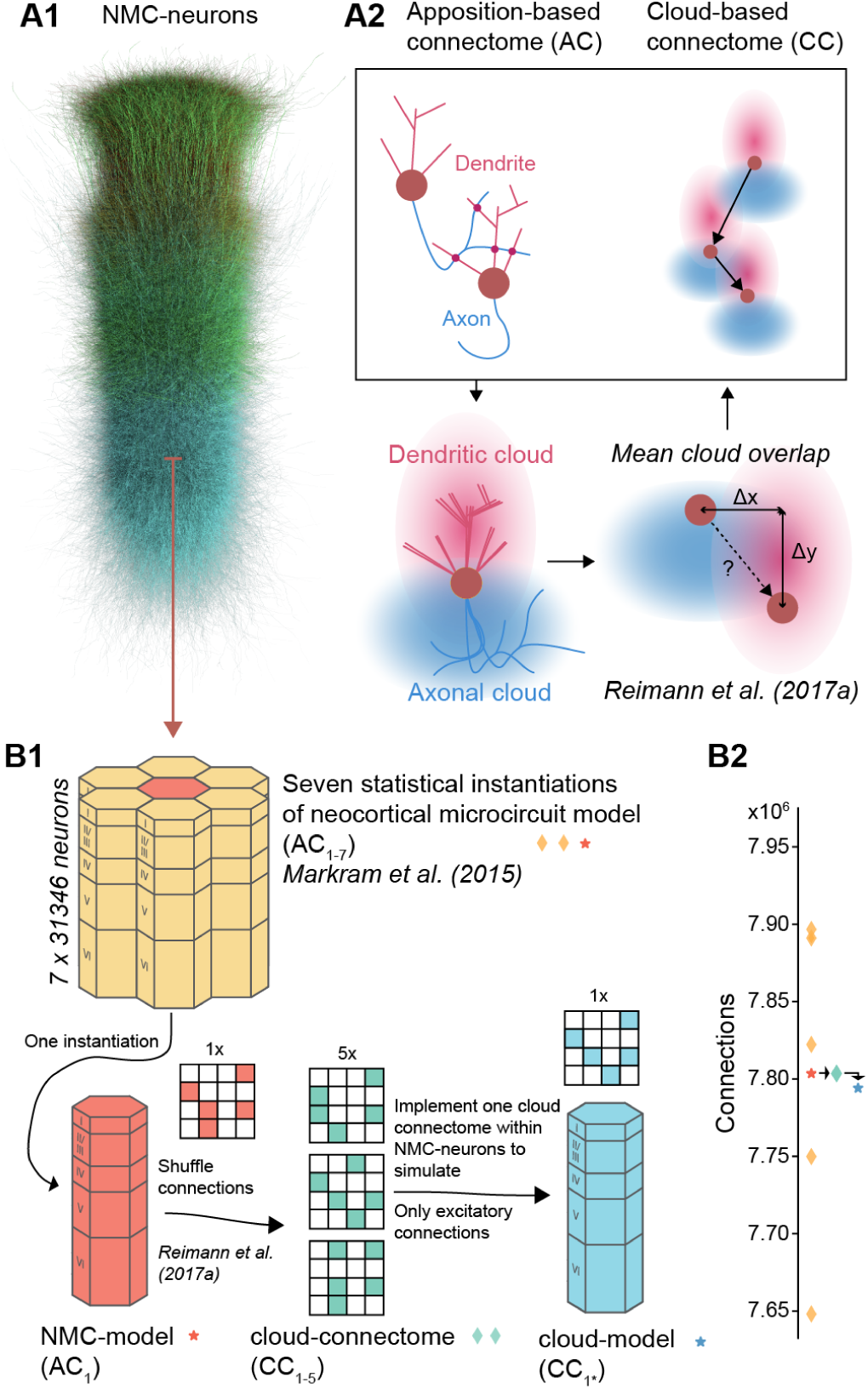
Reducing higher-order network structure in a neocortical microcircuit model. (A1) Visualization of neurons of neocortical microcircuit (NMC-) model. (A2) Deriving synaptic connectivity between neocortical neurons: Connectivity in the NMC-model is based on a touch-based approach that considers appositions of dendrites and axons (Reimann et al., 2015). Connectivity in the control cloud-model considers overlap of average dendritic and axonal clouds (Reimann, Horlemann, et al., 2017). (B1) We computed network properties for seven statistical instantiations of the microcircuit (apposition-based connectomes AC_1−7_, orange diamonds and red star) (Markram et al., 2015), and simulated one of them in this study (the *NMC-model* with connectome AC_1_, red star). Additionally, we studied versions of the model using the existing NMC-neurons, but synaptic connectivity derived according to the cloud-based approach (cloud-connectomes CC_1−5_, green diamonds). We then implemented one of the alternative connectomes within the existing synapses of the NMC-neurons, resulting in the *cloud-model* that we simulated, with connectome C_1*\**_ (blue star). The rewiring of the cloud-model was restricted to excitatory connections. (B2) Number of connections across connectomes. By design, the cloud-connectomes CC_1−5_ have the same number of connections as the NMC-model connectome AC_1_. However, the implemented connectome of the cloud-model CC_1*\**_ has 0.12% fewer total connections due to a mismatch in new connections and available synapses (see Figure 2A).

Here, we utilize a recent finding that first-order connectivity is largely constrained by morphological diversity *between* neuronal types, and higher-order connectivity by morphological diversity *within* neuronal types (Reimann, Horlemann, Ramaswamy, Muller, & Markram, 2017). Both aspects are captured by the NMC-model, leading to a biologically realistic micro-structure (Gal et al., 2017). By connecting neurons according to average axonal and dendritic morphologies (axonal and dendritic *clouds*, Figure 1A2), we create a control circuit (the *cloud-model*), that has very similar first order structure, but reduced higher-order structure. We find that this reduced higher-order structure—caused by disregarding morphological diversity within neuronal types—includes more homogeneous degree distributions, reduced in-degrees at the bottom of layer six, fewer cliques, and decreased small-world topology. When we simulate and compare the electrical activity in the two circuit models, we find that the changes in higher-order connectivity are accompanied by nuanced changes in neuronal firing patterns and reduced topological ordering of pairwise correlations.

Our study introduces a rigorous method to reduce higher-order network structure of a neocortical microcircuit model while leaving first-order structure largely intact and suggests that higher-order network topology of neocortical microcircuitry shapes cortical function.

## RESULTS

### Reducing higher-order network structure in a neocortical microcircuit model

The NMC-model consists of 31,346 neurons belonging to 55 different morphological types (*m-types*) (Figure 1A1). Synaptic connectivity between the neurons was derived by considering appositions of dendrites and axons as potential synapse locations (apposition-based connectome AC_1_, Figure 1A2, left), which were then filtered according to biological constraints (Reimann et al., 2015). While this connectome is merely a null model of connectivity, it matches a large array of biological measurements, both in terms of its *first-order structure* and *higher-order structure*. We define first-order structure as structure of synaptic connectivity that can be described by pairwise statistics, e.g. connection strengths between different m-types and distance-dependent connectivity, and higher-order structure as structure involving more than two neurons (Benson et al., 2016), e.g. common-neighbor bias, over-representation of cliques of all-to-all connected neurons, and also degree distributions. Neuronal and synaptic physiology in the model are equally well constrained (Markram et al., 2015).

To assess the specific role of the higher-order synaptic structure on neuronal activity, we had to reduce the higher-order structure while simultaneously impacting the first-order structure as little as possible. To this end, we used an alternative cloud-based approach to derive synaptic connectivity, which is based on the overlap of average dendritic and axonal shapes of the various m-types (Figure 1A2, bottom, right; *cloud-model*) instead of specific axo-dendritic appositions of individual neurons as in the NMC-model. This approach (see Methods) yields similar properties of first-order structure, such as distinct connection strengths between different m-types (and consequently between layers), distance-dependent connectivity and a bias for reciprocal connections (Reimann, Horlemann, et al., 2017). However, the cloud-model cannot reproduce an experimentally observed bias for connected neocortical neurons to share a common neighbor (Perin et al., 2011; Reimann, Horlemann, et al., 2017), indicating a reduced complexity of its higher-order structure. By comparing electrical activity between the NMC-model and the cloud-model in simulation experiments, we can study the relative impact of first- and higher-order structure on electrical activity.

To build the cloud-model, we first generated alternative cloud-based connectomes for the NMC-neurons (Figure 1B1, red to green), using methods introduced by Reimann, Horlemann, et al. (2017). Briefly, average axon and dendrite shapes were calculated from reconstructions for all m-types. Next, for each combination of neuron types, their axon- and dendrite volumes were convolved (see Figure 1A2) to yield the expected strength of their overlap for all possible relative soma locations. Soma locations of neurons were taken from the NMC-model and used to look up the overlap strengths for all neuron pairs. Connection probabilities for all pairs were then proportional to the square of the overlap and normalized such that the total number of connections for each combination of neuron types matches the connectivity of the NMC-model (Figure 1B2, red asterisk and green diamond).

A neuron-to-neuron connection matrix was then instantiated from the probabilities (the *cloud-connectome*, CC_1−5_) and transplanted into the NMC-neurons, to generate an instance of the *cloud-model* that was identical to the NMC-model in terms of neuronal composition, and morphology and physiology of all individual neurons (Figure 1B1). Similarly, the physiology of individual synapses (strength, kinetics and short-term dynamics) and their locations on dendrites were taken from the NMC-model—we only changed which presynaptic neurons innervated them when implementing the cloud-based connection matrix (Figure 2A). This reassignment of innervation was constrained to select a new innervating neuron only from the same m-type that innervated it in the NMC-model to preserve the m-type-specific synaptic physiology. However, while synaptic physiology is conserved in this approach, the axonal *path length*, and thus the time it takes for an action potential to propagate from the soma to the synapse, is potentially incorrect. While the average path lengths per pre-/postsynaptic m-type combination are conserved, an action potential might potentially arrive earlier or later than is appropriate for the distance between pre- and postsynaptic neuron (see Figure 2A).

**Figure 2.**
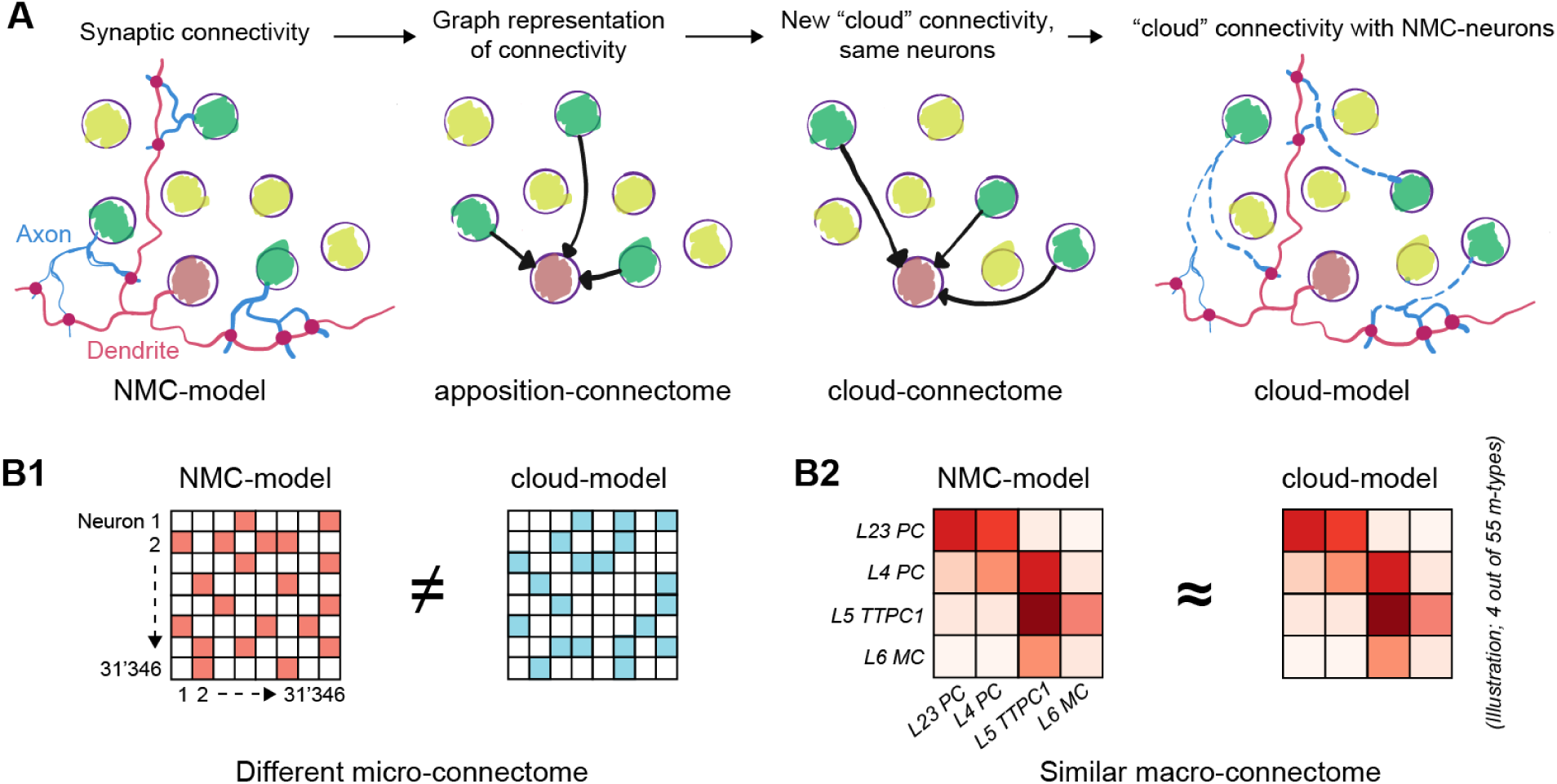
Rewiring synaptic connectivity in a neocortical microcircuit model. (A) The cloud connectome is implemented within the NMC-neurons by using existing synapses of the same connection type, that is identical combinations of pre- and postsynaptic morphological neuron types (*m-type)*. (B) The NMC- and cloud-models have a completely different micro-connectome in terms of connections between individual neurons (B1) but very similar macro-connectome in terms of number of connections between the 55 different m-types in the model (B2).

In the cloud-based connection matrix CC_1_, a small number of neurons receive input from m-types that do not innervate them in the NMC-model. Consequently, a small fraction of connections could not be instantiated within the existing synapses of the NMC-neurons and had to be left out. The loss was minor for excitatory connections (0.12% loss of connections) but posed a significant problem for inhibitory connections. We therefore implemented the cloud-connectome only for excitatory connections and kept inhibitory connectivity in the cloud-model identical to the NMC-model, yielding a hybrid cloud-model connectome CC_1*\**_. Supplementary Figure S1 provides an overview of connection losses in the cloud-model. However, note that the loss of connections is very small compared to the variability in connections between statistical instantiations of the NMC-model (Figure 1B2, orange diamonds), which all have very similar dynamical properties (Markram et al., 2015).

To control for the minor loss of excitatory connectivity and the shuffling of axonal path lengths, we generated an additional control circuit, the NMC*-model. A total of 0.12% of excitatory connections were randomly removed from the NMC-model to match the m-type-specific connection losses in the cloud-model (as in Supplementary Figure S1B). Connections with the same presynaptic excitatory m-type for each neuron were then shuffled and assigned new synapses to account for the scrambling of axonal path lengths in the cloud-model (See Figure 2A). All circuit models and connectomes analyzed in this study are summarized in Table 1.

**Table 1.** Overview of circuit models and connectomes analyzed in this study.

| Name | Description |
| --- | --- |
| **NMC-neurons** | 31,346 neuron models of a neocortical microcircuit (Markram et al., 2015). |
| **AC<sub>1</sub>** | Apposition-based connectome for NMC-neurons (Markram et al., 2015).<sup>a</sup> |
| **NMC-model** | Combination of NMC-neurons and AC<sub>1</sub> (Markram et al., 2015). |
| **AC<sub>2-7</sub>** | Apposition-based connectomes of statistical variants of NMC-model. (Markram et al., 2015).<sup>a</sup> |
| **CC<sub>1-5</sub>** | Cloud-based alternatives to AC<sub>1</sub> (Reimann, Horlemann, et al., 2017). |
| **cloud-model** | Combination of NMC-neurons and CC<sub>1</sub> (excitatory connections) and AC<sub>1</sub> (inhibitory connections). |
| **CC<sub>1\*</sub>** | Connectome of cloud-model (see above). |
| **NMC\*-model** | NMC-model with axon path length shuffle and 0.12% exc. connection loss as in cloud-model. |

<sup>a</sup>mc2\_Column and mc[0, 1, 3-6]\_Column respectively at bbp.epfl.ch/nmc-portal/downloads → AVERAGE.

In summary, our approach ensures that for each neuron in the NMC-model, there is a corresponding neuron in the cloud-model with identical location, morphology, electrophysiology, synaptic physiology, inhibitory innervation and average excitatory innervation patterns. On a larger scale, both models consequently have nearly identical *macro-connectomes* in terms of the number of connections between morphological types (Figure 2B2, and thus also between layers, and between excitatory and inhibitory sub-populations (Supplementary Figure S1C1-3). Only the micro-connectomes defined by connections between individual neurons were changed within tight global constraints (Figure 2B1). An overview of what is conserved between NMC-model, cloud-model, and NMC*-model can be found in Table 2.

**Table 2.**
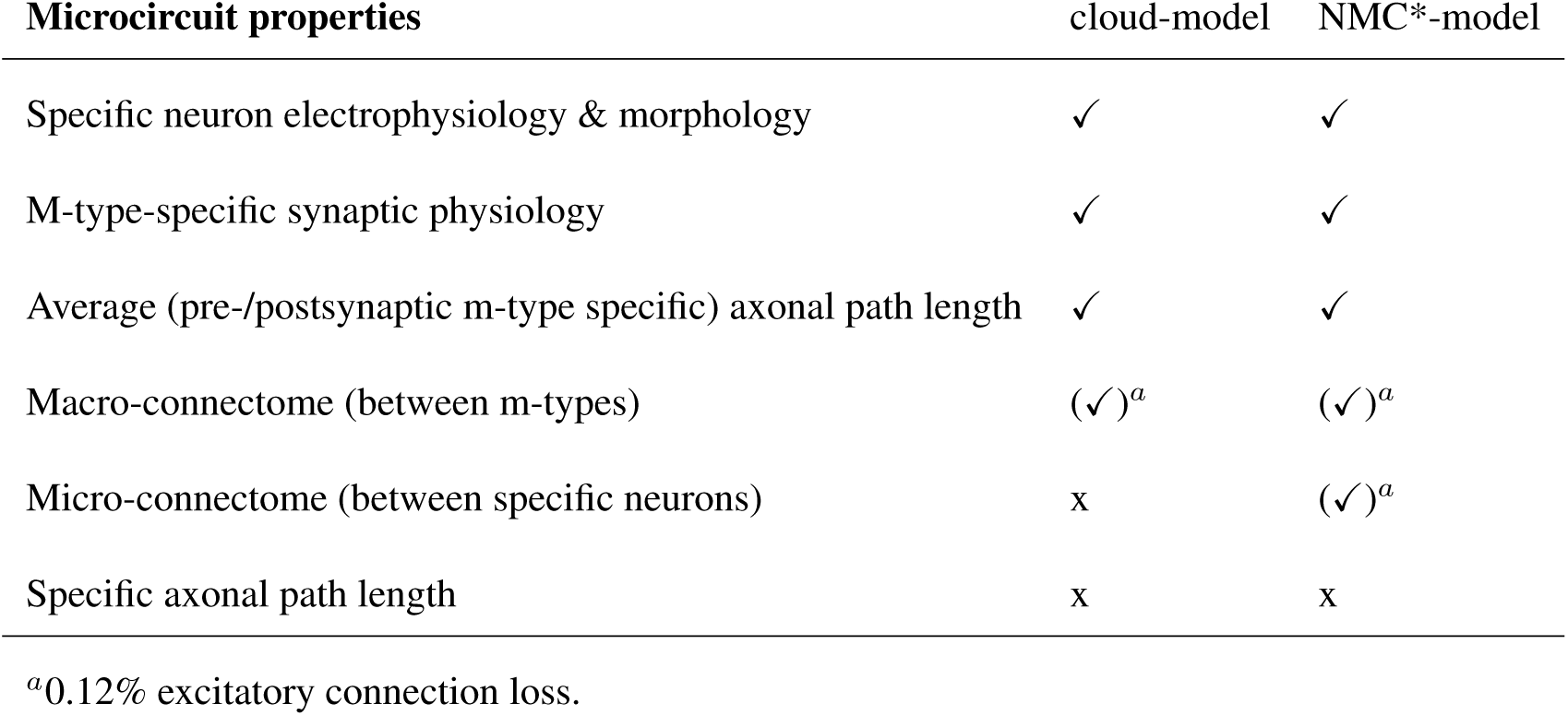
Overview of control model conservation of NMC-model properties.

| Microcircuit properties | cloud-model | NMC*-model |
| --- | --- | --- |
| Specific neuron electrophysiology & morphology | ✓ | ✓ |
| M-type-specific synaptic physiology | ✓ | ✓ |
| Average (pre-/postsynaptic m-type specific) axonal path length | ✓ | ✓ |
| Macro-connectome (between m-types) | (✓)<sup>a</sup> | (✓)<sup>a</sup> |
| Micro-connectome (between specific neurons) | x | (✓)<sup>a</sup> |
| Specific axonal path length | x | x |

<sup>a</sup>0.12% excitatory connection loss.

### Decreased heterogeneity of degree-distributions in cloud-model

The cloud-model has a reduced higher-order structure in terms of a bias for two connected neurons to share common neighbors (*common neighbor bias*) (Reimann, Horlemann, et al., 2017). Other important higher-order characteristics of a network are the in- and out-degree distributions, which directly shape cortical network dynamics (Landau, Egger, Dercksen, Oberlaender, & Sompolinsky, 2016). We can see that both in- and out-degree distributions are more heterogeneous in the NMC-than in the cloud-model (Figure 3A1 and 3A2, blue vs. red), which applies to all layers (Supplementary Figure S2). In the NMC-model, the standard deviation of out-degrees (*σ*_*out*_(AC_1_) = 152, statistical variants: *σ*_*out*_(AC_2−7_) = 150–154 [min–max]) and the standard deviation of in-degrees (*σ*_*in*_(AC_1_) = 176, *σ*_*in*_(AC_2−7_) = 173–179) are higher than in the cloud-model (*σ*_*out*_(CC_1*\**_) = 110, cloud-connectomes: *σ*_*out*_(CC_1−5_) = 101.7–101.8; *σ*_*in*_(CC_1*\**_) = 158, *σ*_*in*_(CC_1−5_) = 158.2–158.4). The discrepancy in *σ*_*out*_ and *σ*_*in*_ of CC_1−5_ and AC_1−7_ is significantly different (*σ*_*out*_: p = 3.0 × 10^−14^, *σ*_*in*_: p = 1.6 × 10^−8^; two-sided t-test). However, the difference between cloud-connectome CC_1_ (cloud-based excitatory *and* inhibitory connections) and the cloud-model CC_1*\**_ (which differs from the NMC-model only in the *excitatory* connections) is small relative to the NMC-model. This suggests that inhibitory connections would only have a minor impact on higher-order structure differences between the NMC- and cloud-models.

**Figure 3.**
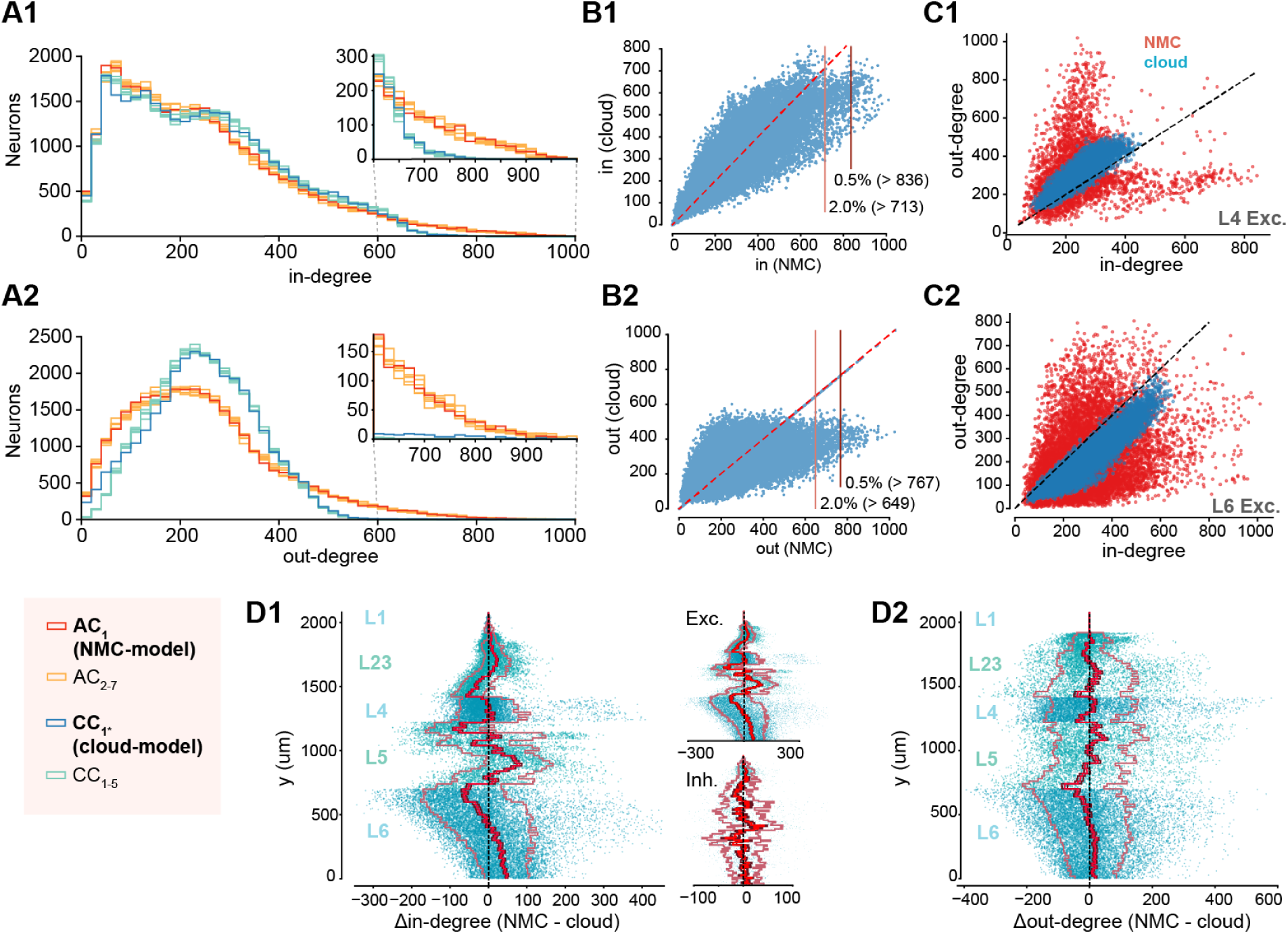
Decreased heterogeneity of degree-distributions in cloud-model. (A1) In-degree distributions of neurons in NMC-model (AC_1_, red), in additional apposition-based connectomes (AC_2−7_, orange), the cloud-model (CC_1*\**_, blue), and the five cloud-connectomes (CC_1−5_, light blue). Insets show the same distributions starting from 600 for easier comparison. (A2) As A1, but for out-degree distributions. (B) Hub neurons in NMC- and cloud-models. (B1) Scatter plot of in-degree in NMC-vs. cloud-model for the same neurons. Horizontal lines indicate 2.0% and 0.5% percentile in NMC-model. (B2) As B1 but for out-degree. (C1) In-vs. out-degree scatter plot for NMC- and cloud-model for excitatory neurons in layer 4. (C2) In-vs. out-degree scatter plot for NMC- and cloud-model for excitatory neurons in layer 6. (D1) Difference in in-degree of neurons between NMC- and cloud-model, across cortical depth. The bright red line indicates the mean across y-bins, the dark red line the standard error of the mean, and the outer, faded line the standard deviation. Insets: for excitatory and inhibitory subpopulations. (D2) As D1, but for out-degree.

The stark difference in connectivity between the NMC- and cloud-models is also reflected by hub neurons, previously defined as the top 0.5% of neurons in terms of in- or out-degree (Gal et al., 2017), which vanish in the cloud-model when using the same cut-off value as in the NMC-model (Figure 3B). The increased heterogeneity of degree distributions in the NMC-model also extends to the correlations between in- and out-degree (Figure 3C1 and 3C2 for excitatory neurons in layers 4 and 6, see Supplementary Figure S3 for all neurons), indicating a stronger specialization into input and output neurons in the NMC-model. We further found a redistribution of connectivity in terms of in- and out-degree from the bottom to the top of layer 6 (Figure 3D1 and 3D2). In summary, the cloud-model has a strongly reduced heterogeneity of connectivity in terms of distributions of in- and out-degrees.

### Decreased small-worldness and fewer directed simplices in cloud-model

The higher-order structure of the NMC-model also manifests itself on a global scale in the form of a small-world network topology (Gal et al., 2017). A network is considered small-world if its global clustering coefficient is considerably larger than that of an Erdös-Rényi (ER) network of the same size and sparseness, while the characteristic path length is roughly equal to that of the ER network. For networks of the same size and sparseness, the ratio of the global clustering coefficient *c* and the characteristic path length *l* provides a measure of relative small-worldness. The clustering coefficient describes the probability of neurons that share a common neighbor to be directly connected, a tendency that has previously been shown to be reduced in the cloud-model (Reimann, Horlemann, et al., 2017). Consequently, we find that the cloud-model has a reduced clustering coefficient (*c* is around 20% larger in the NMC-model than in the cloud-model, Figure 4A2). The characteristic path lengths *l* of NMC- and cloud-models are, however, almost equal to each other (Figure 4A1) and to the one of the equivalent ER network (around 2.15). Both models can therefore be considered small-world networks, although this tendency is significantly stronger in the NMC-model than in the cloud-model (Figure 4A3).

**Figure 4.**
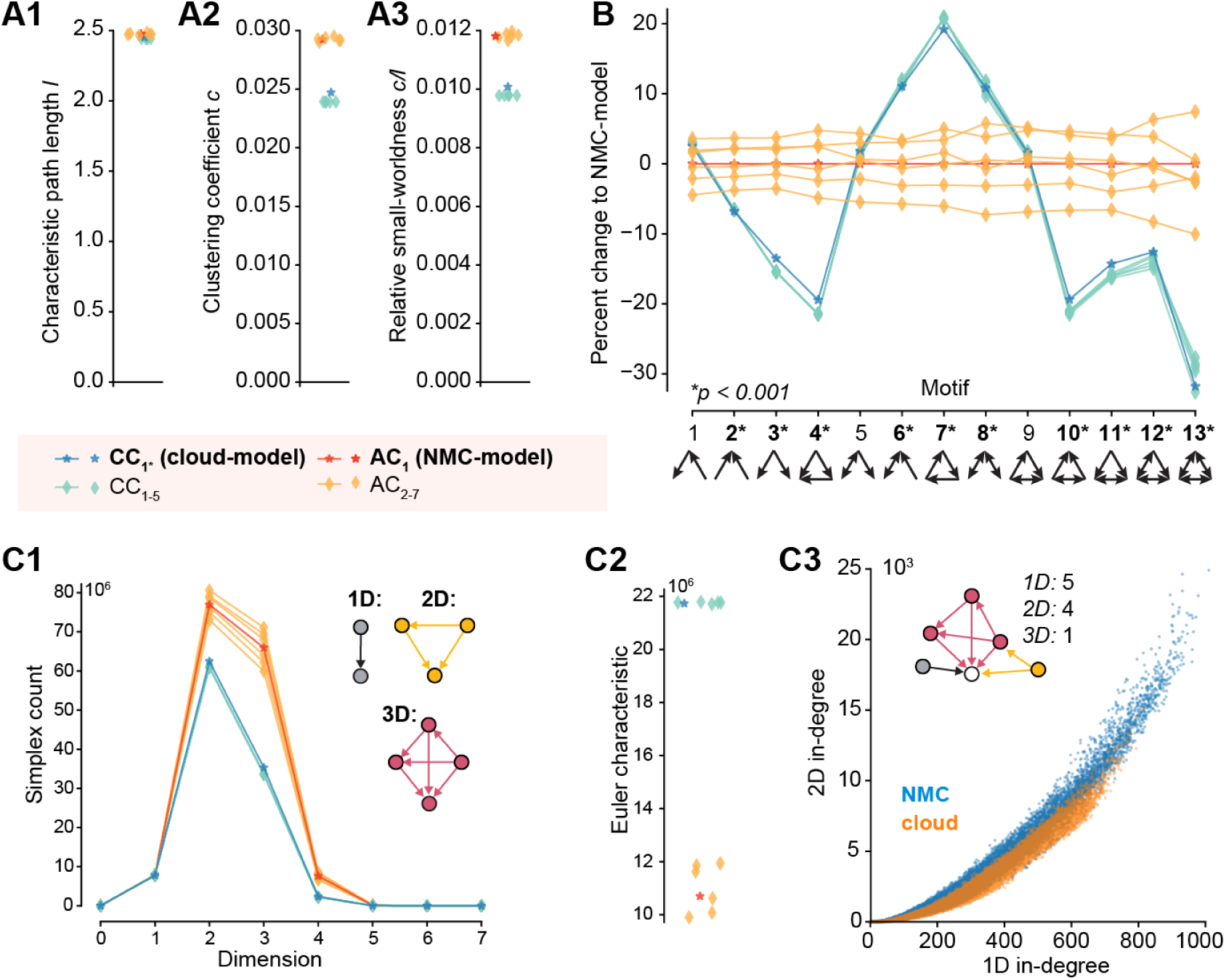
Decreased small-worldness and fewer directed simplices in cloud-model. (A1) Characteristic path length *l* of the different connectomes. (A2) Clustering coefficient *c* of the different connectomes. (A3) Relative small-worldness *c/l* of the different connectomes. (B) Percent change in total number of all triplet motifs from NMC-model (p-values: t-test between AC_1−7_ and CC_1−5_). (C1) Number of directed cliques (simplices) per dimension in the different connectomes (for zoom in see Figure S4A). (C2) Euler characteristic (alternating sum of number of simplices). (A3) Participation at the sink of 1D simplices (standard in-degree) vs. 2D simplices (2D in-degree: the number of simplices a neuron is part of as the sink of a simplex).

As an increased clustering coefficient indicates the tendency to form tightly connected motifs, we next compared the numbers of specific triplet motifs in both models (Figure 4B). Interestingly, the cloud-model has a decrease of motifs with forward transitive connectivity (Figure 4B, motif 4), but an increase in motifs with backwards transitive connectivity, such as cycles (motif 7). We previously showed that the NMC-model contains an abundance of a specific class of motifs called directed simplices (Reimann, Nolte, et al., 2017). These simplices generalize the forward transitive connectivity of motif 4 to motifs of any size. The size of a simplex is then called its dimension, defined as its size minus one. While simplices of the same dimension are present in cloud- and NMC-models (Supplementary Figure S4A), the number of simplices in a given dimension is much higher in the NMC-than in the cloud-model (Figure 4C1). At the same time, the cloud-model is much closer to the NMC-model than simpler control models: in a previously used control that conserves only the distance-dependence of connectivity, but ignores the shapes of axonal and dendritic clouds, we found a more drastic decrease from around 80 million to 40 million 2D-simplices (Reimann, Nolte, et al., 2017), while the cloud-model has more than 60 million 2D-simplices. A distinct Euler Characteristic (Figure 4C2) and distinct Betti-numbers (Supplementary Figure S4B) further illustrate the change in global properties of network topology (see Methods). The increase in simplex numbers in the NMC-model follows from the more heterogeneous degree distributions, as neurons with larger degrees are generally part of more simplices (Figure 4C3).

In summary, the cloud-model—with its disregard for morphological diversity within neuronal types—has reduced small-worldness and reduced numbers of high-dimensional, forward transitive motifs.

### Impact of higher-order structure on spontaneous activity

To study the impact of the higher-order structural differences on emergent activity, we next simulated spontaneous activity in NMC-, cloud- and NMC*-models in an in vivo-like state, in which the NMC-model can reproduce several properties of cortical activity (Markram et al., 2015). We compared firing rates in this in vivo-like state (Figure 5B, [*Ca*^2+^]_*o*_ = 1.25 mM). Interestingly, excitatory firing rates are conserved (Figure 5B2), while inhibitory firing in the NMC*-model is significantly reduced compared to the cloud-model and NMC-model (Figure 5B3). That is, a small decrease of inhibitory firing rates due to the axonal path length shuffle and loss of E-to-I synapses in the NMC*-model is reversed by the reduced higher-order structure of the cloud-model, reaching inhibitory firing rates similar to the NMC-model. This implies that the loss of higher-order structure in the cloud-model leads to a shift towards more inhibition (Figure 5C).

**Figure 5.**
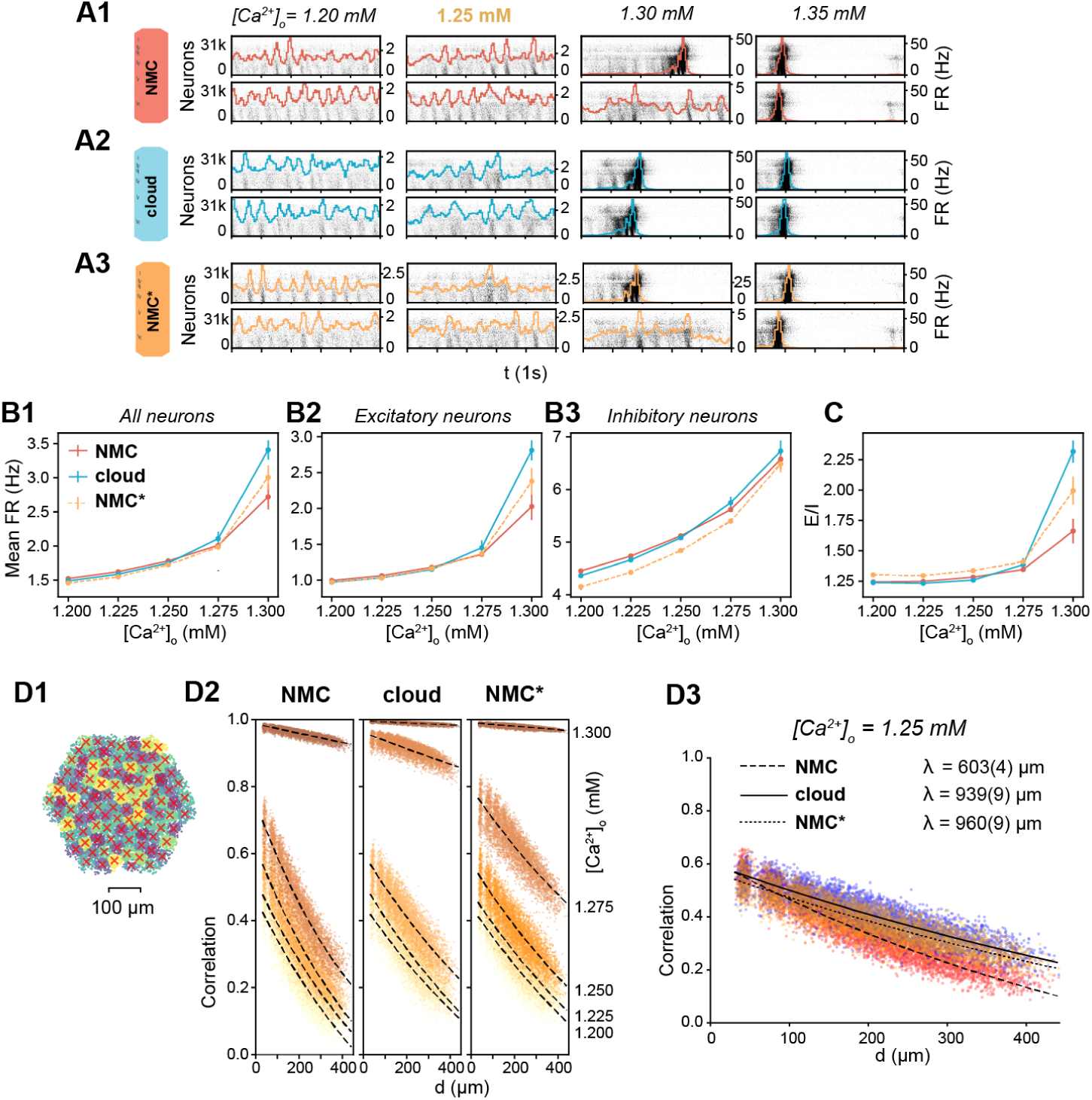
Simulating spontaneous activity in NMC- and cloud-models. (A1) Spontaneous activity of all neurons in the NMC-model for two trials at different levels of [*Ca*^2+^]_*o*_. Each spike is represented by a vertical line, whose position on the y-axis is ordered by soma position in the microcircuit, which was then rasterized. Colored lines depict the population firing rate (Δ*t* = 10 ms). (A2) Spontaneous activity of all neurons in the cloud-model. (A3) Spontaneous activity of all neurons in the control NMC*-model. (B1) Mean firing rate during spontaneous activity for all neurons. Mean of 20 trials of 1000 ms, error bars indicate standard error of the mean. (B2) Mean firing rate during spontaneous activity for excitatory neurons. Mean of 20 trials of 1000 ms, error bars indicate standard error of the mean. (B3) Mean firing rate during spontaneous activity for inhibitory neurons. Mean of 20 trials of 1000 ms, error bars indicate standard error of the mean. (C) Total spike count of excitatory neurons divided by the spike count of inhibitory neurons. Mean of 20 trials of 1000 ms, error bars indicate standard error of the mean. (D1) All neurons in the microcircuit were divided into 100 interlaminar clusters (k-means clustering). The red cross marks the geographic center of all neurons for each cluster. (D2) Correlation-coefficients between combined firing rate histograms of all neurons of each combination of clusters vs. distance between clusters. From [*Ca*^2+^]_*o*_ = 1.200 mM (bottom fitted curve, bright yellow dots) to [*Ca*^2+^]_*o*_ = 1.300 mM (top fitted curve, dark brown dots). The fitted line indicates an exponential fit *e*^*-d/λ*^ + *c*. (D3) As in D2, but at [*Ca*^2+^]_*o*_ = 1.25 mM. The fitted line indicates an exponential fit *e*^*-d/λ*^ + *c*, parentheses indicate the standard error of the fit.

However, the functional excitation-inhibition balance also depends on extracellular calcium levels ([*Ca*^2+^]_*o*_), which differentially modulates the release probability of excitatory and inhibitory synapses (Markram et al., 2015). In the NMC-model, at low [*Ca*^2+^]_*o*_, the network is in an asynchronous state of activity with lower firing rates (Figure 5A1, [*Ca*^2+^]_*o*_ = 1.2 mM, 1.25 mM). At high [*Ca*^2+^]_*o*_, the circuit is in a non-biological synchronous state of activity with spontaneous network bursts and higher firing rates (Figure 5A1, [*Ca*^2+^]_*o*_ = 1.3 mM, 1.35 mM). At [*Ca*^2+^]_*o*_ = 1.25 mM, just before the transition from the asynchronous to synchronous state, activity in the microcircuit is in the in vivo-like state described above.

The cloud-model appears to transition at lower [*Ca*^2+^]_*o*_ than the NMC-model and NMC*-model, with more rapidly increasing firing rates for higher [*Ca*^2+^]_*o*_ (Figure 5B1–3), an indication that effective excitation is stronger in the cloud-model than in NMC- and NMC*-models. [*Ca*^2+^]_*o*_ also regulates distance-dependent correlation coefficients of spiking activity between neurons (Figure 5D1) (Markram et al., 2015). At the transition from asynchronous to synchronous activity, correlation coefficients rapidly increase (Figure 5D2). We can see that at [*Ca*^2+^]_*o*_ = 1.25 mM, correlations between NMC- and NMC*-model are very similar, but the cloud-model is in fact slightly ahead, with higher correlation coefficients (Figure 5D2). At that level of [*Ca*^2+^]_*o*_, correlations also drop slightly faster with distance in the cloud-than in the NMC-model, however, this is fully explained by the non-topological changes controlled for in the NMC*-model (Figure 5D3).

We thus conclude that the higher-order network structure has a superficial impact on emergent population dynamics during spontaneous activity, such as a small increase in inhibitory firing rates, and paradoxically, a small increase in effective global excitation.

### Impact of higher-order structure on evoked activity

Spontaneous activity in the NMC-model is variable and chaotic, yet, thalamic stimuli can evoke highly reliable responses (Nolte, Reimann, King, Markram, & Muller, 2019). To study the impact of higher-order structure on this evoked activity, we next stimulated the NMC- and cloud-models with thalamic input (Figure 6A1, see Methods). Similar to spontaneous activity, the response of the circuits to the input depended on [*Ca*^2+^]_*o*_ (Figure 6A2-4). As we established that the NMC- and cloud-models have a slightly different transition between dynamic states, we compared evoked activity for up to five [*Ca*^2+^]_*o*_ values around the in vivo-like state. All three models exhibited similar fluctuations of the overall firing rate at various [*Ca*^2+^]_*o*_ levels, responding robustly with brief increases in firing to periods of correlated thalamic input (Figure 6A2-4). As before, mean firing rates are very similar between the models, but the cloud-model has increased excitation, especially for larger [*Ca*^2+^]_*o*_ (Figure 6C1, Supplementary Figure S5A1).

**Figure 6.**
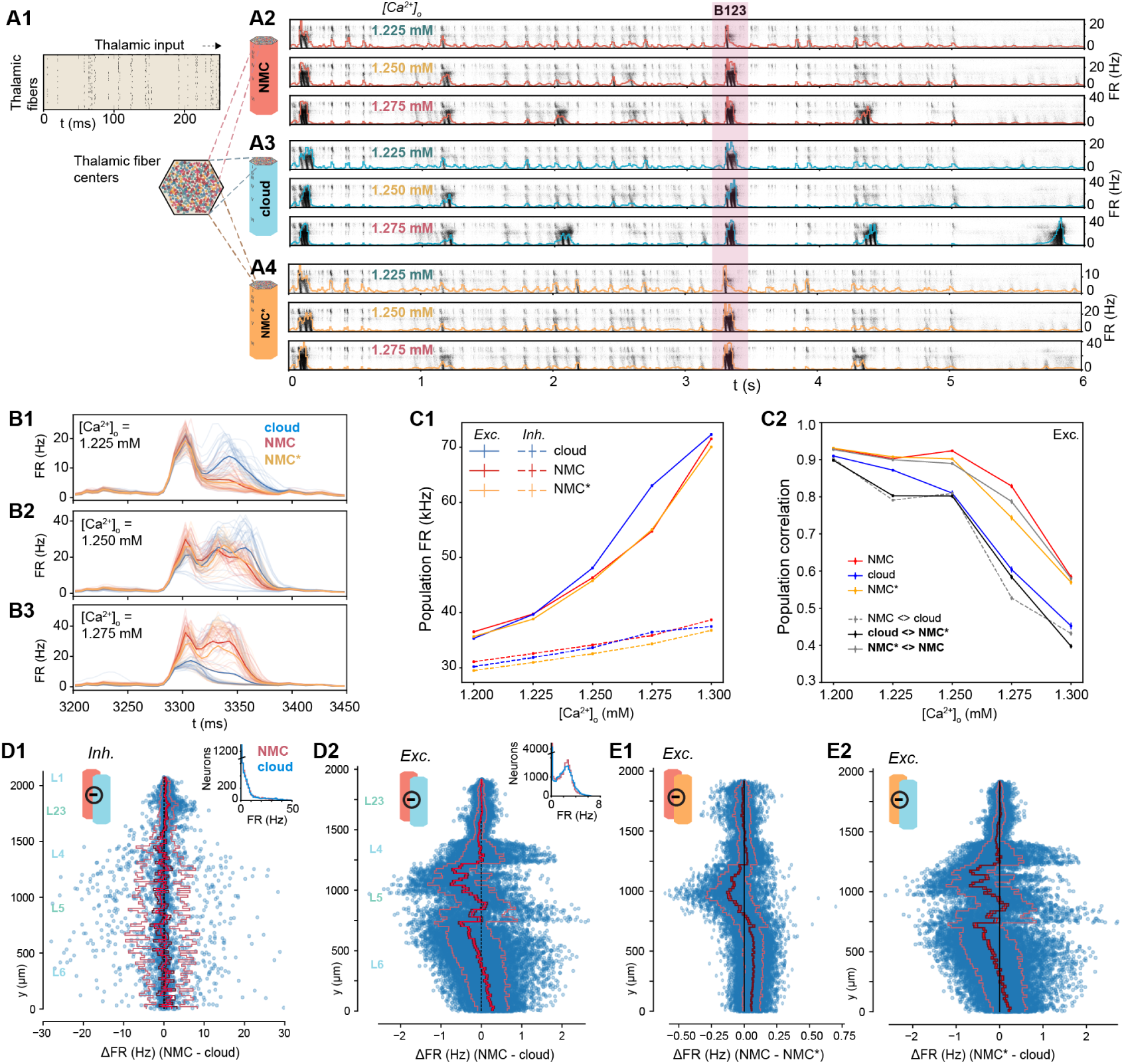
Simulating evoked activity in NMC- and cloud-models. (A1) Illustration of the first 250 ms of the thalamic input stimulating the microcircuit from *t* = 0 to 5 s. (A2) Evoked activity for all neurons in the NMC-model in response to input from A1 at three different [*Ca*^2+^]_*o*_-levels. (A3) Evoked activity for all neurons in the cloud-model at three different [*Ca*^2+^]_*o*_-levels. (A4) Evoked activity for all neurons in the NMC*-model at three different [*Ca*^2+^]_*o*_-levels. (B1) Time-dependent population firing rate for NMC-model (red), cloud-model (blue), and NMC*-model (orange) for a 250 ms time period during the evoked activity (shaded red area in A234) at [*Ca*^2+^]_*o*_ = 1.225 mM. Faint lines: population means for all 30 trials. Thick line: mean of all 30 trials. Bins size: Δ*t* = 5 ms. (B2) As B1, but for [*Ca*^2+^]_*o*_ = 1.25 mM. (B3) As B1 and B2, but for [*Ca*^2+^]_*o*_ = 1.275 mM. (C1) Mean population firing rate of excitatory and inhibitory subpopulations during evoked activity at different [*Ca*^2+^]_*o*_-levels. Error bars indicate standard error of the mean over 30 trials. Note that error bars are smaller than linewidth. (C2) Average correlation coefficient between excitatory population PSTHs (Δt = 5 ms) for 30 trials. Mean of 30 × (30 − 1)*/*2 = 435 combinations for same model, and mean of 30 *×* (30 + 1)*/*2 = 465 combinations between models. (D1) Difference in firing rate of inhibitory neurons during evoked activity at [*Ca*^2+^]_*o*_ = 1.25 mM. Blue dots indicate values for individual neurons, ordered along their soma positions with respect to the y-axis (cortical depth). Lines indicate mean (bright red), standard-error (dark red), and standard deviation. Inset: Distribution of all mean firing rates. (D2) As D1, but for excitatory neurons. (E1) As D2, but for NMC-model and NMC*-model. (E2) As D1, but for NMC*-model and cloud-model.

To gain a more detailed understanding of the difference, we next looked at the changes in firing rates for individual neurons. Excitatory firing rates varied according to the change in in-degree across cortical depth (Figure 6D2 vs. Figure 3D1). The mean change directly reflects the change in in-degree shown in Figure 2D1—which is not surprising as any change in in-degree in the cloud-model is restricted to excitatory connections. While both excitatory and inhibitory neurons respond to the change in in-degree individually (Supplementary Figure S5B), the systematic shift in in-degree only emerges for excitatory, but not inhibitory, neurons (see insets in Figure 3D1).

Time-dependent response patterns differ between models and [*Ca*^2+^]_*o*_-levels (Figure 6B1-3), with distinct patterns and increased trial-by-trial variability in the cloud-model when compared to activity in NMC-models and NMC*-models. To quantify this difference, we calculated the correlation coefficients of the peri-stimulus time histograms (PSTHs) between individual trials of the same model and different models (Figure 6C2, Supplementary Figure S5A2). The correlation between trials of the same model—that is, the reliability of the population response—generally decreased with increasing[*Ca*^2+^]_*o*_-level, although for the NMC- and NMC*-models it remained over 0.9 until 1.25 mM and was significantly higher than for the cloud-model at all levels. Correlations between different models were highest between NMC- and NMC*-models, further reinforcing that the slight loss of excitatory connections alone does not explain the observed changes in response patterns of the cloud-model.

We have previously shown that the in-degree also influences the spike-time reliability (*r*_spike_) of individual neurons in response to repeated trials of a thalamic stimulus. (Nolte et al., 2019). While there is overall little change in *r*_spike_ going from NMC- to cloud-model (Supplementary Figure S6ABC), we observed a drop in reliability near the top of layer 5, which also displays a large in-degree reduction in the cloud-model. Further, as neurons with reduced in-degree towards the bottom of layer 6 spike less, they also become less reliable (Supplementary Figure S6BD). Similar to the firing rate, the change in spike-time reliability is clearly correlated with the change in in-degree (Supplementary Figure S6E).

### Impact on ordering of correlations in simplices

We have shown that the cloud-model has a reduced bias for forward-transitive triplet motifs (Figure 4B), resulting in a reduced number of directed simplices (Figure 4C1). We have further demonstrated that this leads to changes in the activity patterns of the circuit, specifically a reduced reliability of the population response to thalamic input. It has been shown that reliable responses are linked to increased correlations of synaptic inputs (Nolte et al., 2019; Wang, Spencer, Fellous, & Sejnowski, 2010), and we previously observed that such input correlations can be generated by directed simplex motifs with stronger correlations found in larger simplices (Reimann, Nolte, et al., 2017).

We therefore analyzed the structure of spike-time correlations of pairs of neurons in simplices across models (Figure 7A). As before (see Figure 5D), the overall strengths of correlations were comparable, though slightly higher in the cloud-model at identical [*Ca*^2+^]_*o*_-levels. This effect can be explained by the shift along the spectrum from asynchronous to synchronous activity (see above). However, we observed a qualitative difference in the local structure of correlations within a simplex. As a directed structure, each simplex can be uniquely sorted from the *source* neuron, with only outgoing connections to all other neurons in the simplex, to the *sink* neuron, with only incoming connections (Figure 7B4). While the correlations increased with simplex size for all three models (Figure 7B1–3), for the NMC- and NMC*-models they strongly depended on the location of neurons in the simplex, with highest correlations for the pair at the sink and lowest correlations for the pair at the source. Conversely, in the cloud-model this difference was reduced and disappeared for the largest simplex sizes.

**Figure 7.**
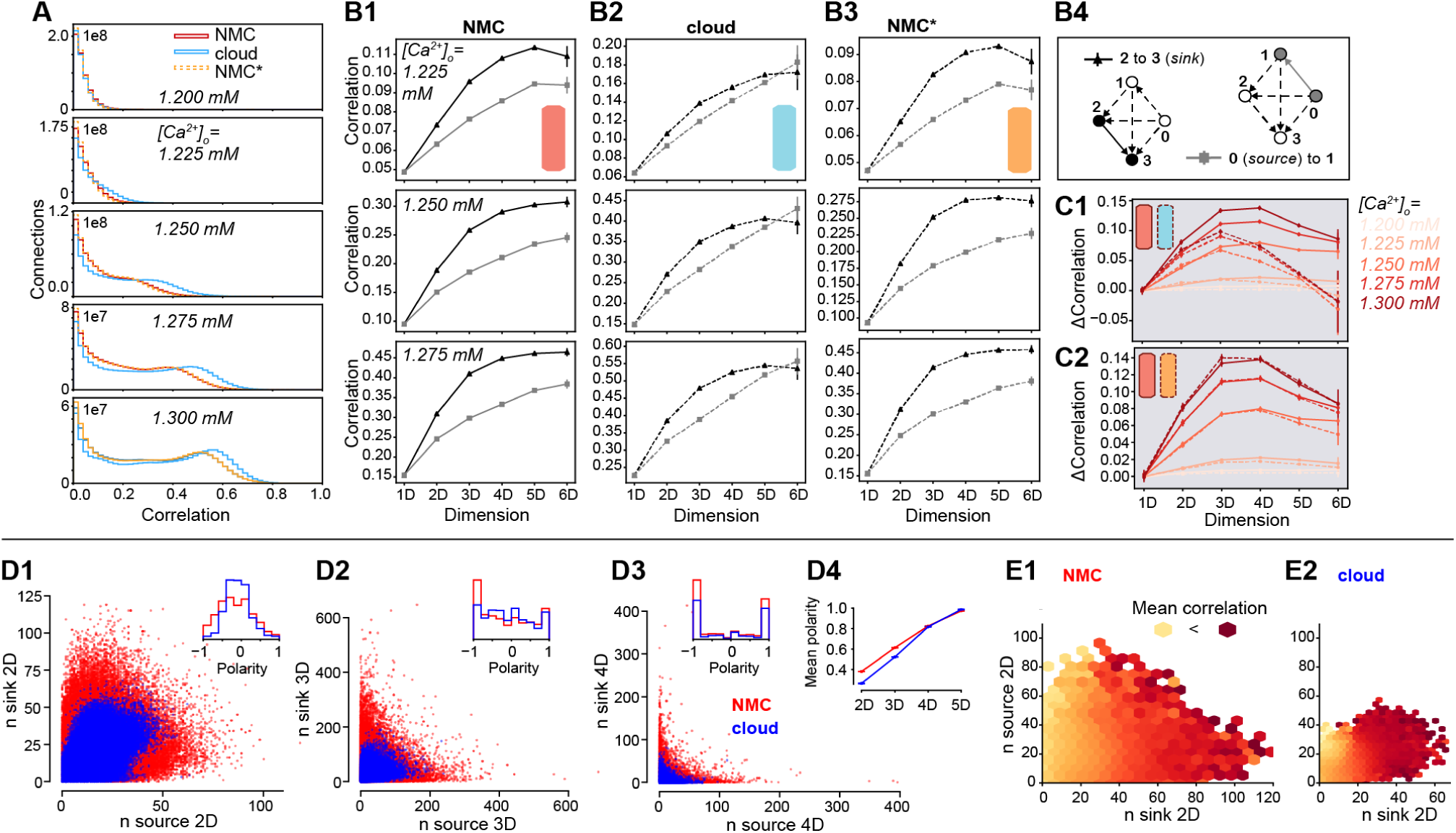
Simplices and correlations in NMC- and cloud-models. (A1) Correlation coefficients of the firing rates (Δ*t* = 20 ms) of all connected pairs of active neurons in the microcircuit for NMC-model, cloud-model, and NMC*-model, during the 30 trials of the thalamic stimulus. (B1) Average correlation coefficient of a connected pair of neurons in a (maximal) simplex of a certain dimension, depending on the position of the pair in the simplex; for the NMC-model, for three different [*Ca*^2+^]_*o*_-levels. Black triangles indicate the average correlation for a pair of neurons at the *sink* of a simplex, grey squares the average correlation for a pair at the *source* of a simplex. (B2) As B1, but for cloud-model. (B3) As B1, but for NMC*-model. (C1) Difference in average correlation coefficient for connections of neurons at the sink and the source of a (maximal) simplex. Solid lines: NMC-model; dashed lines: cloud-model. (C2) Difference in average correlation coefficient for connections of neurons at the sink and the source of a (maximal) simplex. Solid lines: NMC-model; dashed lines: NMC*-model. (D1) Participation of connections in the source vs. sink of (non-maximal) 2D-simplices in NMC- and cloud-models (constrained to connections at the center of layers 4, 5, 6). Inset: Polarity is defined as 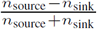. (D2) As D1, but for 3D-simplices. (D3) As D1, but for 4D-simplices. (D4) Mean polarity for each dimension, defined as mean of 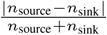. (E1) Mean correlation given the participation of a connection in (non-maximal) 2D-simplices at the source and sink, from bright yellow (low correlation) to dark red (high correlation); in NMC-model, constrained to connections at the center of layers 4, 5, 6. (E2) As D1, but for the cloud-model.

In other words, correlations in simplices have a hierarchical organization that leads to a polarization between input (source) neurons and output (sink) neurons that is diminished in the cloud-model. This is consistent with the earlier finding that higher correlations of in- and out-degrees of the cloud-model prevent a polarization into input- and output-neurons on a structural level (see Figure 3C). In fact, this effect extends from the in- and out-degree of individual neurons to an equivalent measure of simplex participation of connections. For each connection we count the number of simplices in which it forms the first connection at the source, equivalent to the out-degree, and the number of simplices in which it forms the last connection at the sink, equivalent to the in-degree. As before, we found higher degrees with lower correlations between in- and out-degree in the NMC-model (Figure 7D1–3). We calculated the *polarity* of a connection as the relative difference between its in- and out-degree with respect to simplices of a given dimension. We found that its absolute value increased with simplex dimension and was overall higher in the NMC-model, indicating a stronger structural polarization (Figure 7D4).

Taken together, we hypothesize that each simplex provides correlated input to neuron pairs at its sink, which in turn leads to correlated firing of that pair. While the effect of a single simplex on a pair may be negligible, strong structural polarization into inputs and outputs suggests that such pairs are likely to participate as the sink of an unexpectedly high number of simplices, thus dramatically increasing the size of the effect. The stronger polarization of the NMC-model further increases this effect. This hypothesis predicts that the correlation of a neuron pair is determined by the in-degree of its connection and is unaffected by its out-degree. We therefore quantified how the spike-correlations of neuron pairs depend on these measures (Figure 7E). Indeed, we found a strong dependence on the in-but not the out-degree for both models.

## DISCUSSION

We introduced a method to reduce the higher-order structure of synaptic connectivity in a neocortical microcircuit model, based on a previously published control connectome with a reduced common-neighbor bias (Reimann, Horlemann, et al., 2017)—the so-called cloud-model. In this cloud-model, excitatory synaptic connectivity between neurons was derived from average morphologies rather than appositions of axons and dendrites of individual neurons (Figure 1). We showed that the reduction in higher-order structure in the cloud-model includes more homogeneous in- and out-degree distributions, the disappearance of hub neurons (Figure 3), less all-to-all directed cliques of neurons and reduced small-worldness (Figure 4). Spontaneous and evoked dynamics in the NMC-model and cloud-model are superficially very similar (Figures 5-6), a result that is not surprising given the conserved first-order structure, including conserved interlaminar connectivity, structural excitation-inhibition balance and distance-dependent connectivity. However, some properties of neuronal activity changed. Spike counts of individual neurons changed according to in-degree and population responses were shifted towards more excitation, with less reliable population responses (Figure 6). Most importantly, a hierarchical dependence of correlation strength on the position of a pair of neurons in directed motifs (simplices) was weaker in the cloud-model than in the NMC-model (Figure 7). We have demonstrated that this was a result of the more homogeneous in- and out-degree distributions leading to a reduced polarization into input- and output-neurons, and consequently reduced participation in simplices.

Consistent with previous definitions of higher-order network structure (Benson et al., 2016), our definition includes degree distributions of neurons, which are not constrained by the pairwise statistics of neurons alone. Nevertheless, reducing higher-order network structure while keeping degree distributions fixed is possible, and such higher-order structure can have an impact on network dynamics (Ritchie, Berthouze, House, & Kiss, 2014). To better understand the respective impact of changes in in- and out-degrees on one side, and clustering and high-dimensional motifs on the other side, it will be necessary to create a more refined control model that conserves degree distributions on top of first-order structure.

The comparison between NMC- and cloud-models comes with several caveats. Distance-dependent connectivity in the cloud-model is not perfectly preserved for all m-type combinations (Reimann, Horlemann, et al., 2017). Distance-dependance is however much better conserved than in previous control models that disregarded the average shapes of m-types, resulting in many fewer simplices (Reimann, Nolte, et al., 2017). The cloud-model further neglects to reduce inhibitory higher-order structure. However, the structural similarity between cloud-connectomes (changed excitatory and inhibitory connectivity) and the cloud-model (changed excitatory connectivity only) suggests that the contribution of inhibitory connections to higher-order network structure in the NMC-model is negligible. This could potentially be due to an underestimation of inhibitory higher-order structure in the NMC-model, either due to insufficient biological data constraining the connectivity, or because inhibitory structure might only emerge through plasticity (Vogels, Sprekeler, Zenke, Clopath, & Gerstner, 2011).

Indeed, this brings us to the most important caveat: the NMC-model is a statistical reconstruction of a prototypical microcircuit from sparse data. While it captures a high level of detail of synaptic connectivity (Gal et al., 2017), with strong constraints on the space of connectivity that can be explored by structural plasticity (Reimann, Horlemann, et al., 2017), the model has not learned to respond to specific stimuli or perform certain computations. A comparison to numbers of simplices observed in in vitro slice experiments shows that the number of simplices in the NMC-model is likely underestimated by an order of magnitude (Reimann, Nolte, et al., 2017). This suggests that a large fraction of biological higher-order structure is not captured by the NMC-model. This leads to at least two interesting questions to be explored in the future: 1) what is the functional impact of that additional higher-order structure on network dynamics and 2) given a biologically plausible model of structural plasticity, could the network reach similar higher-order structure starting from both NMC- and cloud-models?

Our method of separating first and higher-order structure need not only be applied to statistical connectome models, but might also be useful for the interpretation of future dense reconstructions of brain tissue using electron microscopy (Kasthuri et al., 2015). Once a volume large enough to contain several neurons of different m-types can be reconstructed, comparing the cloud connectivity of average reconstructed neurons to the actual biological connectivity could serve as a powerful control to interpret the structure of synaptic connectivity. In particular, it can help quantify how much structure emerges from plasticity mechanisms (Zhang, Zhang, & Stepanyants, 2018), and how much is determined by other factors, such as neuronal morphologies (Reimann, Horlemann, et al., 2017).

In summary, we introduced a method for investigating the functional impact of higher-order network structure of neocortical microcircuitry. Going beyond the common neighbor bias described by Reimann, Horlemann, et al. (2017), our analysis demonstrates just how many higher-order structural properties are constrained by neuronal diversity within neuronal types. Our comparison between the two models suggests that the higher-order network structure of cortical synaptic connectivity impacts emergent dynamics and might be a non-negligible component of cortical function.

## METHODS

### Circuit models

#### Neocortical microcircuit model (NMC-model)

Methods are based on a previously published model of a neocortical microcircuit of the somatosensory cortex of the two week-old rat, here called the *NMC-model* (Markram et al., 2015). Synaptic connectivity (with apposition-based connectome/adjacency matrix AC_1_) between 31,346 neurons belonging to 55 different morphological types (m-types) was derived algorithmically starting from the appositions of dendrites and axons, and then taking into account further biological constraints such as number of synapses per connection and bouton densities (Reimann et al., 2015). Neuronal activity in the NMC-model was then simulated in the NEURON simulation environment (www.neuron.yale.edu/neuron/). Detailed information about the circuit, NEURON models and the seven connectomes of the different statistical instantiations of the NMC-model analyzed in this study are available at bbp.epfl.ch/nmc-portal/ (Ramaswamy et al., 2015). Simulations and analysis were performed on an HPE SGI 8600 supercomputer (BlueBrain5).

#### cloud-connectome

Synaptic connectivity based on average morphologies (with cloud-based connectome/adjacency matrix CC) was computed with methods previously described by Reimann, Horlemann, et al. (2017). In brief, for each m-type *m*_*i*_ out of the 55 m-types, we computed 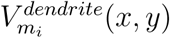 and 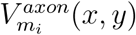, the mean dendrite and axon density of each m-type, based on 10 reconstructed morphologies per m-type, with a resolution of 2*µm* × 2*µm*. Next, we computed for all combinations of m-types the convolution of axon and dendrite densities 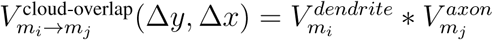. This yielded a measure of the expected strength of the overlap of axon and dendrite for pairs of neurons at all potential relative soma positions. We then looked up this value for all pairs of neurons of a given combination of m-types *m*_*i*_ → *m*_*j*_, based on their locations in the NMC-model and formed a matrix of overlap strengths 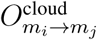 (stored in a table). We next applied a transfer function *Õ* = *O*^2^, which was chosen to conserve distance-dependent connectivity from the NMC-model for most m-type combinations (Reimann, Horlemann, et al., 2017). We then normalized the matrix 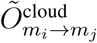 to yield a matrix of connection probabilities such that the expected number of connected pairs equals the number of pairs in the NMC-model:

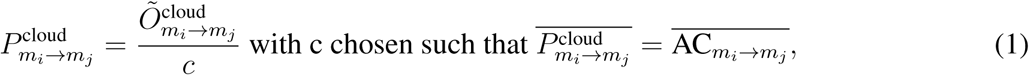

where 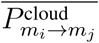 is the average connection probability for *m*_*i*_ → *m*_*j*_ in the cloud-model, and 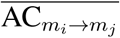 is the average connection probability for *m*_*i*_ → *m*_*j*_ in the NMC-model. The cloud-based adjacency matrix CC was then randomly generated from the normalized connection probabilities for all 55 × 55 *m*_*i*_ → *m*_*j*_ combinations.

Five example connectomes CC_1−5_ for each of the seven NMC-connectomes AC_1−7_ are available at bbp.epfl.ch/nmc-portal/downloads → AVERAGE (Reimann, Horlemann, et al., 2017).

#### cloud-model

We implemented one of the cloud-model connectomes(CC_1_) within the existing neurons of the NMC-model, using pre-existing synapses from the NMC-model. To keep physiological properties such as mean number of synapses per connection conserved, we rewired connections by changing the *source* of *netCon* in NEURON for all synapses in a connection to a new presynaptic neuron according to CC_1_, and we constrained this rewiring to connections with the same presynaptic m-type in the cloud-as in the NMC-model. If there were less connections of a *m*_*i*_ → *m*_*j*_ combination than required by CC_1_, we duplicated connections and their synapses. As some neurons receive input in CC_1_ from neurons that they did not receive input in in AC_1_, some connections could not be implemented. This was a particular problem for inhibitory connections, and we therefore only implemented CC_1_ for excitatory neurons. This resulted in a connectivity matrix CC_1*\**_ that uses CC_1_ for excitatory connections (with a 0.12% loss of connections), and conserved connectivity AC_1_ for inhibitory connections.

#### NMC*-model

To ensure that any changes in emergent activity were not due to the 0.12% of missing connections, or to a shuffling of path lengths (shuffled delays of action potential propagation from soma to synapse), we created a control circuit in which we randomly removed exactly the same number of connections per m-type combination (0.12%) from the NMC-model as could not be implemented in the cloud-model, i.e.:

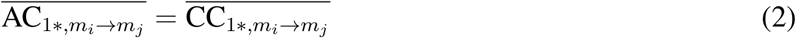

We then shuffled connections with the same presynaptic m-type for each postsynaptic neuron (only for excitatory neurons), keeping the connectivity matrix identical but the locations of synapses randomized as in the cloud-model (and consequently incorrect axonal path lengths).

### Simulation

#### Spontaneous activity

Simulation methods are identical to methods described by Markram et al. (2015): To simulate spontaneous activity, neurons were injected with a depolarizing somatic current, with currents expressed as a percent of first spike threshold for each neuron (100% current used). The *U*_*SE*_ parameter for synaptic transmission of inhibitory and excitatory synapses was then differentially modulated by changing the extracellular calcium concentration [*Ca*^2+^]_*o*_. At [*Ca*^2+^]_*o*_ = 1.25 mM, the circuit was in an in vivo-like state of asynchronous activity with a global balance of excitation and inhibition. We simulated 20 trials of spontaneous activity (two seconds) in the NMC-model, cloud-model, and NMC-model_cloud-control_ at five different [*Ca*^2+^]_*o*_ concentrations around the in vivo-like state at [*Ca*^2+^]_*o*_ = 1.25 mM. We further added two trials of two-second duration at other [*Ca*^2+^]_*o*_ concentrations to illustrate the transition from asynchronous to synchronous activity. The first second of activity was discarded, as the circuit does not reach a resting state until the second second.

#### Evoked activity

We simulated spontaneous activity for seven seconds, as described above. After one second (at *t* = 0 ms, as we discard the first second) we apply a thalamic stimulus through synapses of 310 VPM fibers that innervate the microcircuit. The stimulus lasts five seconds (*t* = 0 to 5 s) and is identical to a previously described stimulus (Nolte et al., 2019; Reimann, Nolte, et al., 2017) based on in vivo thalamic recordings to whisker deflection (Bale, Ince, Santagata, & Petersen, 2015). We simulated 30 trials of the same stimulus in the NMC-model, cloud-model and NMC-model_cloud-control_ at five different [*Ca*^2+^]_*o*_ concentrations around the in vivo-like state at [*Ca*^2+^]_*o*_ = 1.25 mM.

### Analysis

#### In-degree

Number of presynaptic connections a neuron forms with other neurons in the microcircuit.

#### Out-degree

Number of postsynaptic connections a neuron forms with other neurons in the microcircuit.

#### Simplices

A *simplex* is a clique of all-to-all connected neurons. Methods and definitions were adapted from Reimann, Nolte, et al. (2017). In brief, if *G* = (*V, E*) is a directed graph, where *V* is a set of vertices (neurons) and *E* a set of ordered pairs of vertices (directed connections between neurons), then its *directed nD-simplices* for *n* ≥ 1 are (*n* + 1)-tuples (*v*_0_, …, *v*_*n*_) of vertices such that for each 0 ≤ *i* < *j* ≤ *n*, there is an edge in *G* directed from *v*_*i*_ to *v*_*j*_. Neuron 0 (the vertex *v*_0_), the *source* of the simplex (*v*_0_, …, *v*_*n*_), receives no input from within the simplex, but innervates all neurons in the simplex (there is an edge directed from *v*_0_ to *v*_*i*_ for all 0 < *i* ≤ *n*). Neuron 1 (*v*_1_), receives input from Neuron 0, and innervates Neurons 2 (*v*_1_) to *n* (*v*_*n*_), and so forth. Neuron *n*, the *sink*, receives input from all neurons in the simplex, but does not innervate any (there is an edge directed from *v*_0_ to *v*_*i*_ for all 0 < *i* ≤ *n*). See Figure 2E1 for an illustration. Note that reciprocal connections are counted separately: an *n*-simplex in *G* is defined by the (ordered) sequence (*v*_0_, …, *v*_*n*_), but not by the underlying set of vertices (neurons). For instance (*v*_1_, *v*_2_, *v*_3_) and (*v*_2_, *v*_1_, *v*_3_) are distinct 2*D*-simplices with the same neurons. We computed simplices using *flagser* (https://github.com/luetge/flagser).

Note that to avoid redundancy, the average correlation in Figures 6ABC is for *maximal* simplices, simplices that are not part of any higher dimensional simplices (Reimann, Nolte, et al., 2017), whereas all other figures show all simplices, including ones that are fully contained within simplices of a higher-dimension.

#### Higher-order in-degree

We define the *N* D-in-degree as the number of *N* D-simplices a neuron is the sink of. For 1D-simplices (a pair of connected neurons), this is simply the in-degree.

#### Simplex participation of pairs of neurons

We define as the *N* D-participation of connections at the source or the sink of a simplex how many *N* D-simplices a connection is part of as the source (neurons 0 and 1) or at the sink (neurons *N* - 1 and *N*).

#### Betti numbers

In brief, Betti numbers describe the number of cavities or “holes” formed by the simplices in each dimension. Betti numbers were computed using *flagser* (https://github.com/luetge/flagser). Detailed methods are as previously described by Reimann, Nolte, et al. (2017).

#### Euler characteristic

The alternating sum of the number of simplices in each dimension (and of non-zero Betti-Numbers).

#### Small-worldness

Methods are as defined by Gal et al. (2017) and were computed using the Brain Connectivity Toolbox (Rubinov & Sporns, 2010). In brief, we first computed the characteristic path length of the network, defined as the mean shortest path length averaged across all pairs of mutually reachable neurons:

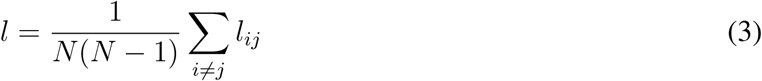

where *l*_*ij*_ denotes the length of the shortest path from neuron *i* to *j*. We next defined the network-wide clustering coefficient as:

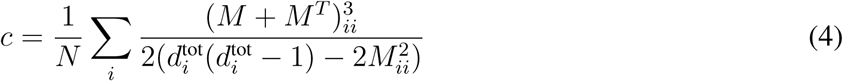

where *M* is the binary connection matrix and 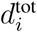 the combined in- and out-degree of each neuron *i*. Thus, *c* = 0 indicates that there are no common neighbors, and *c* = 1 indicates that all neighbors are mutually connected. The ratio of *c/l* gives indication about the small-worldness of the network. We showed previously that the NMC-model has a small-world topology by comparing it to different control models (Gal et al., 2017). A smaller value of *c/l* for the cloud-than NMC-model thus shows that small-worldness decreases.

#### Triplet motif counting

Triplet motifs were counted using the netsci Python library (Gal, Perin, Markram, London, & Segev, 2019) available at https://github.com/gialdetti/netsci.

#### Firing rate

We defined the firing rate (FR) as the average number of spikes in a time bin of size Δ*t*, divided by Δ*t*.

#### Spike-time reliability

Spike-time reliability was quantified with a correlation-based measure (Schreiber, Fellous, Whitmer, Tiesinga, & Sejnowski, 2003). The spike times of each neuron *n* in each trial *k* (*K* = 30 trials) were convolved with a Gaussian kernel of width *σ*_*S*_ = 5*ms* to result in filtered signals *s*(*n, k*, ; *t*) for each neuron *n* and each trial *k* (Δ*t*_*S*_ = 0.5*ms*). For each neuron *n*, the spike-timing reliability is defined as the mean inner product between all pairs of signals divided by their magnitude:

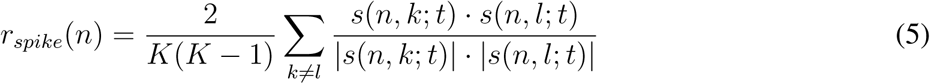

Computation of spike-time reliability is identical to a previous study (Nolte et al., 2019).

#### Correlation coefficients

We computed peristimulus time histograms (PSTHs) for each neuron *i* to the 30 trials of the thalamic stimulus (with a bin size Δ*t*=20 ms), and next computed the normalized covariance matrix of the PSTHs of all neurons:

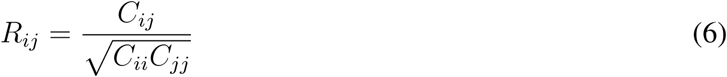

*C*_*ij*_ is the covariance of PSTHs of neurons *i* and *j*. The analysis is replicating a previous analysis by Reimann, Nolte, et al. (2017). Population based correlation coefficients used aggregated PSTHs of the population of neurons (either subpopulation for spatial correlations, or whole circuit for correlations between trials and circuits).

## SUPPORTING INFORMATION

Supporting information includes Supplementary Figures S1-S6.

## ACKNOWLEDGMENTS

We thank the Blue Brain team for developing and maintaining the microcircuit model and computational infrastructure. We thank Kathryn Hess, Idan Segev and Eilif Muller for helpful discussions. We thank Taylor H. Newton for help in editing the manuscript. This study was supported by funding to the Blue Brain Project, a research center of the École Polytechnique Fédérale de Lausanne, from the Swiss governments ETH Board of the Swiss Federal Institutes of Technology. E.G. was supported by the Drahi Family Foundation to Idan Segev, and a grant from the EU Horizon 2020 program (720270, Human Brain Project).

## AUTHOR CONTRIBUTIONS

Conceptualization, M.N., M.R., H.M.; Methodology, M.N., M.R.; Software, M.N., M.R.; Validation, M.N., M.R.; Investigation, M.N., E.G.; Visualization, M.N.; Writing – Original Draft, M.N.; Writing – Review & Editing, M.N., M.R.; Supervision, M.R., H.M.

## SUPPLEMENTARY FIGURES (S1 TO S6)

**Figure S1.**
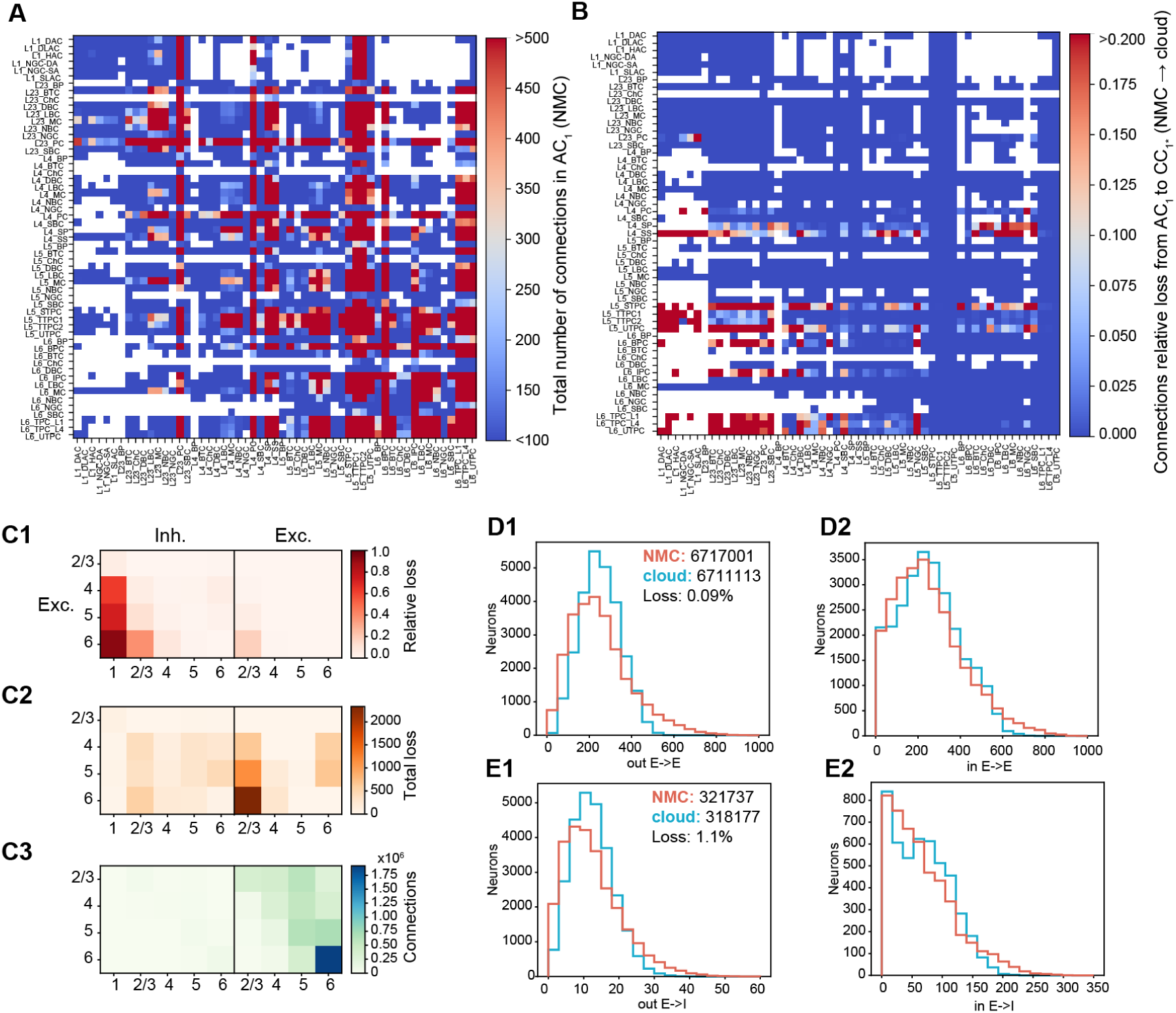
Macro-connectome of NMC- and cloud-models. (A) Total number of connections between m-types in NMC-model. (B) Relative loss of connections from NMC- to cloud-model between each combination of pre- and postsynaptic m-types. (C1) Relative loss of connections from NMC- to cloud-model between layers and excitatory and inhibitory neurons. (C2) Total loss of connections from NMC-to cloud-model between layers and excitatory and inhibitory neurons. (C3) Total number of connections in NMC-model between layers and excitatory and inhibitory neurons. (D1) Out-degree of excitatory neurons counting only excitatory (*E*) connections. Numbers describe total numbers of E-to-E connections in NMC- and cloud-models. (D2) In-degree of excitatory neurons counting only excitatory connections. (E1) Out-degree of excitatory neurons counting only connections formed with postsynaptic inhibitory (*I*) neurons. Numbers describe total numbers of E-to-I connections in NMC- and cloud-models. (E2) In-degree of inhibitory neurons counting only excitatory connections.

**Figure S2.**
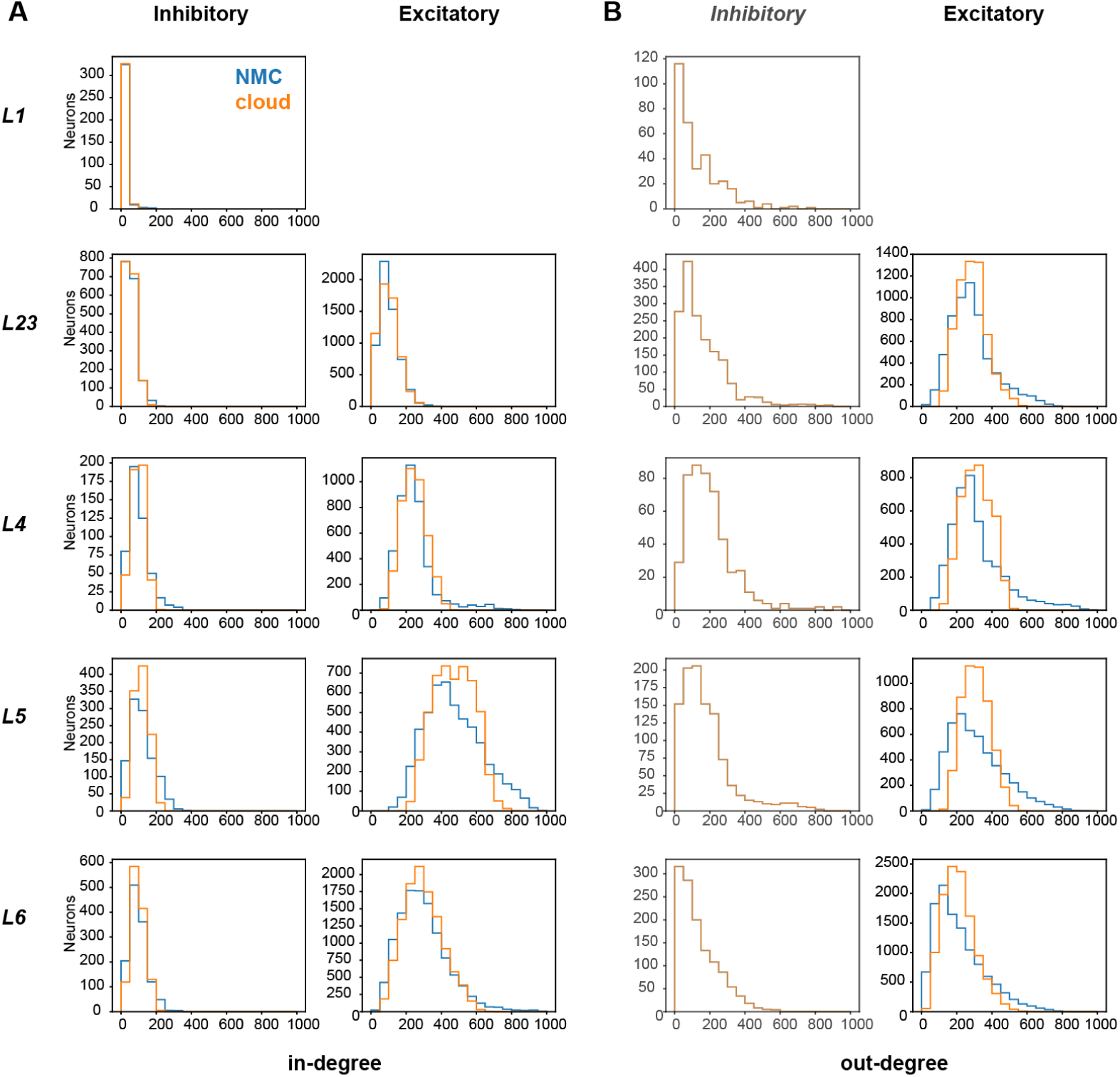
In- and out-degrees across layers. (A) In-degrees of neurons in NMC- and cloud-models, by layer and excitatory/inhibitory sub-type. (B) Out-degrees of neurons in NMC- and cloud-models, by layer and excitatory/inhibitory sub-type.

**Figure S3.**
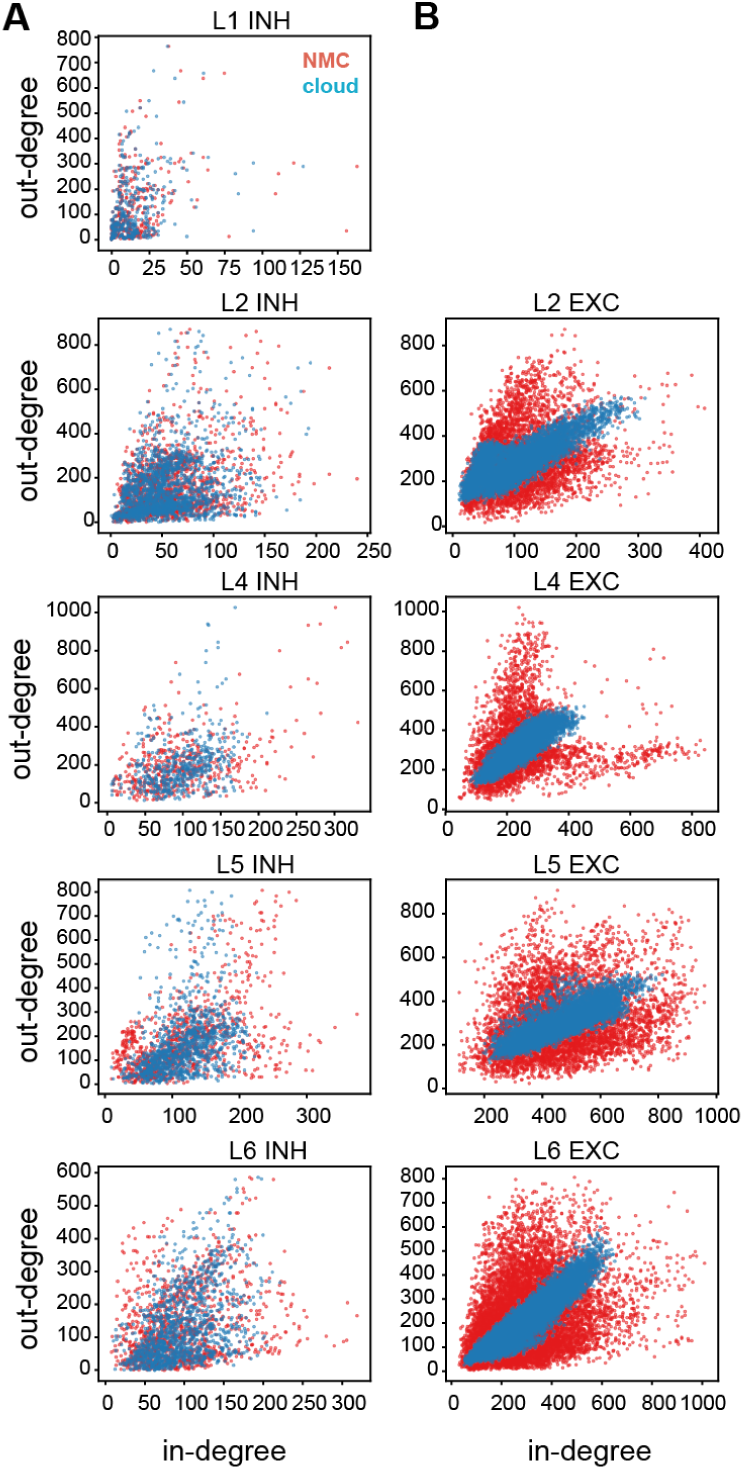
In- and out-degree correlations. (A) Correlations between in- and out-degree for all inhibitory neurons in NMC- and cloud-models, sorted by layer. (B) Correlations between in- and out-degree for all excitatory neurons in NMC- and cloud-models, sorted by layer.

**Figure S4.**
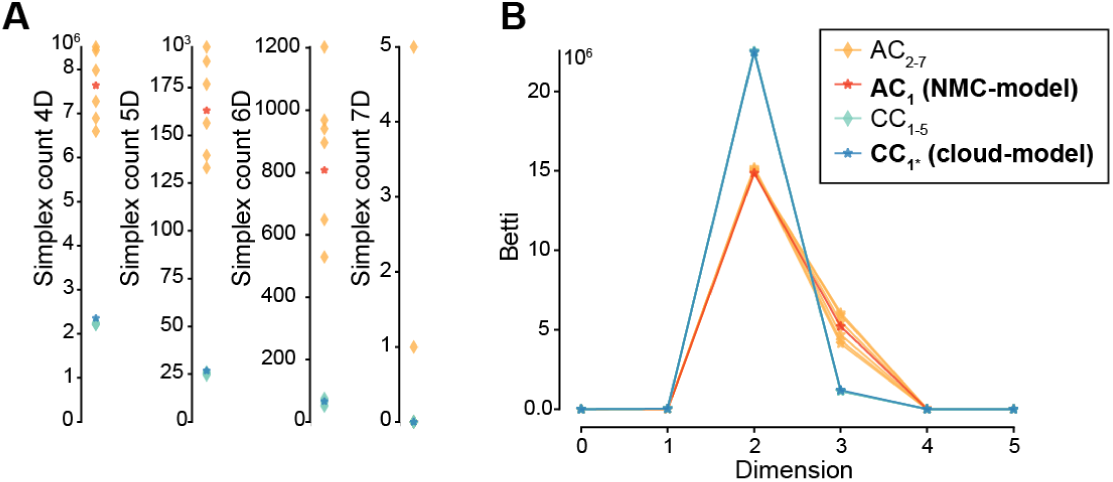
High-dimensional simplices and Betti-numbers. (A) Number of simplices of dimensions 4, 5, 6 and 7 across connectomes. Same legend as B. (B) Betti-numbers across connectomes.

**Figure S5.**
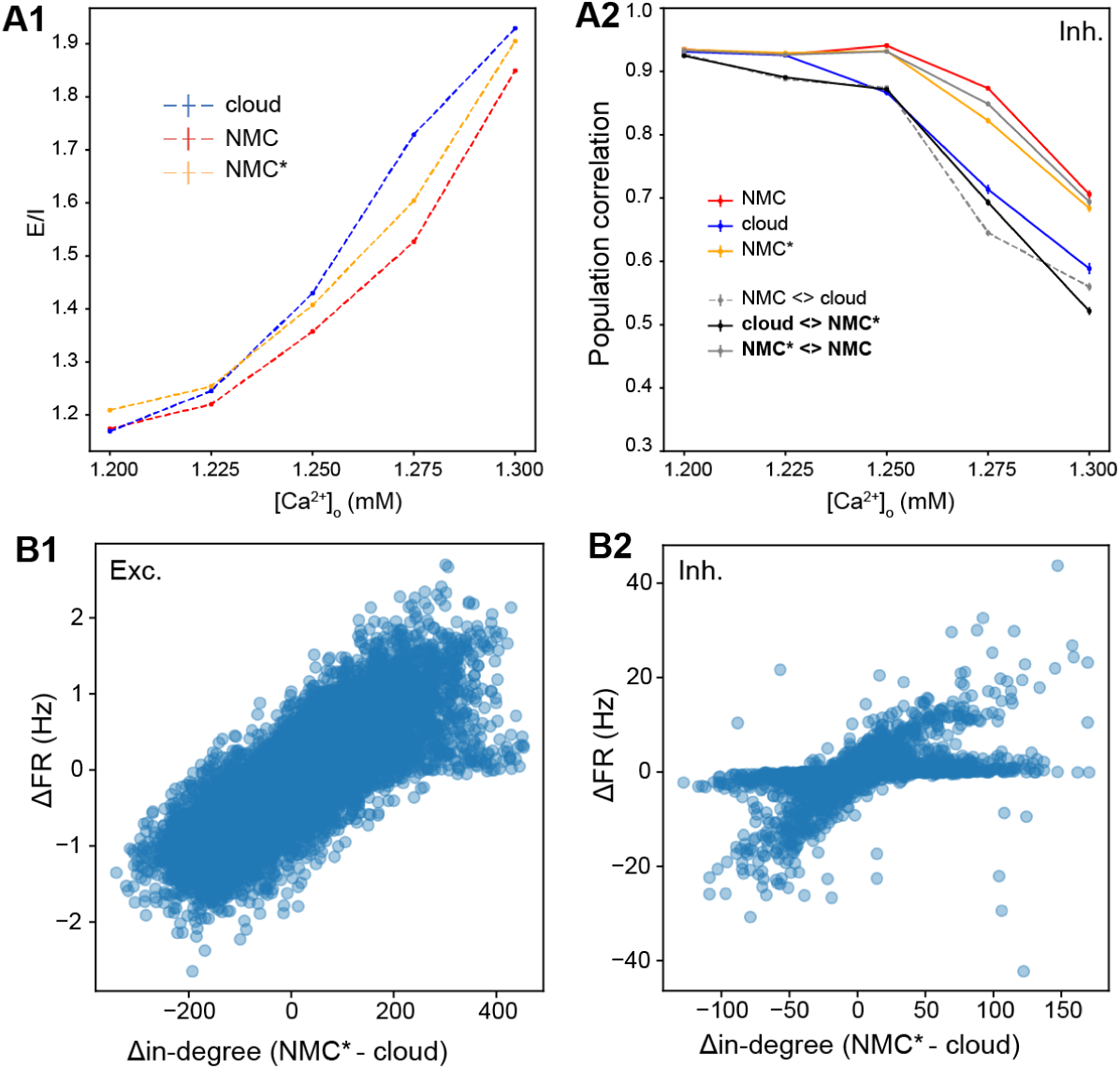
Firing rates and in-degree. (A1) Total spike count of excitatory neurons divided by the spike count of inhibitory neurons. Mean of 30 trials of 6000 ms, error bars indicate standard error of the mean. (A2) Average correlation coefficient between inhibitory population PSTHs (Δt = 5 ms) for 30 trials. Mean of 30 *×* (30 − 1)*/*2 = 435 combinations for same model, and mean of 30 × (30 + 1)*/*2 = 465 combinations between models. (B1) Difference in firing rate during six seconds of evoked activity between NMC*-model and cloud-model vs. difference in in-degree. Blue dots indicate values for individual excitatory neurons. (B2) As B1, but for inhibitory neurons.

**Figure S6.**
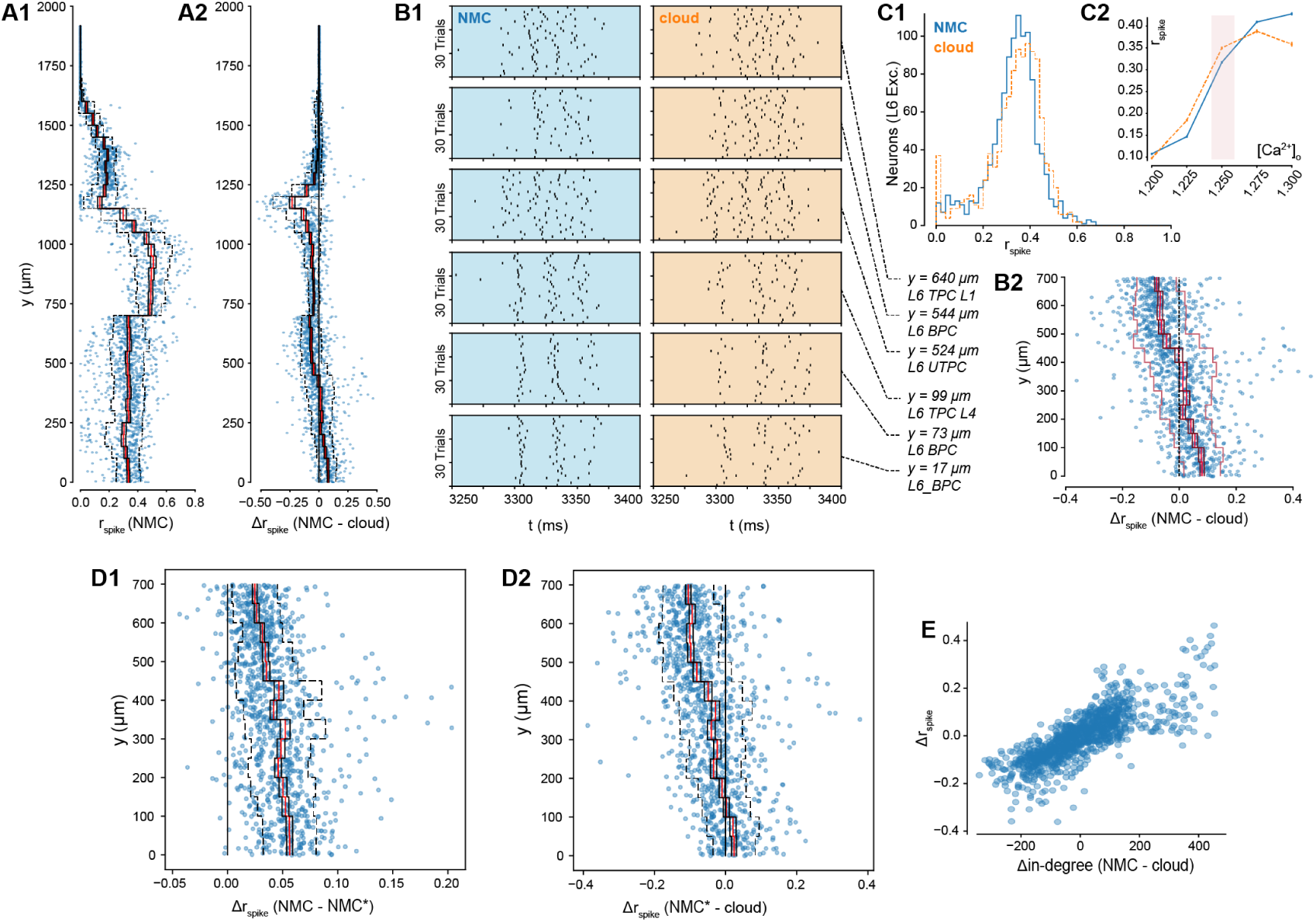
Spike-time reliability. (A1) Spike-time reliability *r*_spike_ of neurons across layers in NMC-model, at [*Ca*^2+^]_*o*_ = 1.25 mM. (A2) Difference in *r*_spike_ of neurons across layers in NMC- and cloud-models at [*Ca*^2+^]_*o*_ = 1.25 mM. (B1) Raster plot of six selected layer six (L6) excitatory neurons during 30 trials of evoked activity, at [*Ca*^2+^]_*o*_ = 1.25 mM. (B2) Difference in spike-time reliability of L6 excitatory neurons during evoked activity at [*Ca*^2+^]_*o*_ = 1.25 mM between NMC-model and cloud-model. Blue dots indicate values for individual neurons, ordered along their soma positions with respect to the y-axis (cortical depth). Lines indicate mean (bright red), standard-error (black), and standard deviation (dashed). (C1) Spike-time reliability *r*_spike_ values for individual L6 excitatory neurons for NMC- and cloud-models at [*Ca*^2+^]_*o*_ = 1.25 mM. (C2) Mean of C1 at various [*Ca*^2+^]_*o*_ levels. (D1) As B2, but between NMC- and NMC*-models. (D2) As B2, but between NMC*- and cloud-models. (E) Difference in Firing rate vs. spike-time reliability between NMC- and cloud-models at [*Ca*^2+^]_*o*_ = 1.25 mM.

## REFERENCES

Bale, M. R., Ince, R. A. A., Santagata, G., & Petersen, R. S. (2015). Efficient population coding of naturalistic whisker motion in the ventro-posterior medial thalamus based on precise spike timing. Front Neural Circuits, 9. Retrieved 2016-11-04, from http://www.ncbi.nlm.nih.gov/pmc/articles/PMC4585317/ doi: 10.3389/fncir.2015.00050

Benson, A. R., Gleich, D. F., & Leskovec, J. (2016). Higher-order organization of complex networks. Science, 353(6295), 163–166. Retrieved 2019-05-11, from https://science.sciencemag.org/content/353/6295/163 doi: 10.1126/science.aad9029

Braitenberg, V. (1978). Cell Assemblies in the Cerebral Cortex. In Theoretical Approaches to Complex Systems (pp. 171–188). Springer, Berlin, Heidelberg. Retrieved 2017-04-24, from https://link.springer.com/chapter/10.1007/978-3-642-93083-6_9 (DOI: 10.1007/978-3-642-93083-69)

Feldmeyer, D., Lubke, J., Silver, R. A., & Sakmann, B. (2002). Synaptic connections between layer 4 spiny neurone-layer 2/3 pyramidal cell pairs in juvenile rat barrel cortex: physiology and anatomy of interlaminar signalling within a cortical column. The Journal of Physiology, 538(3), 803–822. Retrieved 2013-08-13, from http://www.jphysiol.org/cgi/doi/10.1113/jphysiol.2001.012959 doi: 10.1113/jphysiol.2001.012959

Fino, E., & Yuste, R. (2011). Dense Inhibitory Connectivity in Neocortex. Neuron, 69(6), 1188–1203. Retrieved 2014-02-24, from http://linkinghub.elsevier.com/retrieve/pii/S0896627311001231 doi: 10.1016/j.neuron.2011.02.025

Gal, E., London, M., Globerson, A., Ramaswamy, S., Reimann, M. W., Muller, E., … Segev, I. (2017). Rich cell-type-specific network topology in neocortical microcircuitry. Nature Neuroscience, advance online publication. Retrieved 2017-06-06, from https://www.nature.com/neuro/journal/vaop/ncurrent/full/nn.4576.html doi: 10.1038/nn.4576

Gal, E., Perin, R., Markram, H., London, M., & Segev, I. (2019). Neuron Geometry Underlies a Universal Local Architecture in Neuronal Networks. bioRxiv, 656058. Retrieved 2019-05-31, from https://www.biorxiv.org/content/10.1101/656058v1 doi: 10.1101/656058

Hebb, D. (1949). The organization of behaviour. New York: Wiley & Sons.

Holmgren, C., Harkany, T., Svennenfors, B., & Zilberter, Y. (2003). Pyramidal cell communication within local networks in layer 2/3 of rat neocortex. The Journal of Physiology, 551(1), 139–153. Retrieved 2013-08-15, from http://www.jphysiol.org/cgi/doi/10.1113/jphysiol.2003.044784 doi: 10.1113/jphysiol.2003.044784

Jiang, X., Shen, S., Cadwell, C. R., Berens, P., Sinz, F., Ecker, A. S., … Tolias, A. S. (2015). Principles of connectivity among morphologically defined cell types in adult neocortex. Science, 350(6264), aac9462–aac9462. Retrieved 2017-05-03, from http://www.sciencemag.org/cgi/doi/10.1126/science.aac9462 doi: 10.1126/science.aac9462

Kasthuri, N., Hayworth, K. J., Berger, D. R., Schalek, R. L., Conchello, J. A., Knowles-Barley, S., … Lichtman, J. W. (2015). Saturated Reconstruction of a Volume of Neocortex. Cell, 162(3), 648–661. Retrieved 2019-03-19, from https://www.cell.com/cell/abstract/S0092-8674(15)00824-7 doi: 10.1016/j.cell.2015.06.054

Knoblauch, A., Palm, G., & Sommer, F. T. (2009). Memory Capacities for Synaptic and Structural Plasticity. Neural Computation, 22(2), 289–341. Retrieved 2017-04-26, from http://www.mitpressjournals.org/doi/full/10.1162/neco.2009.08-07-588 doi: 10.1162/neco.2009.08-07-588

Landau, I. D., Egger, R., Dercksen, V. J., Oberlaender, M., & Sompolinsky, H. (2016). The Impact of Structural Heterogeneity on Excitation-Inhibition Balance in Cortical Networks. Neuron, 92(5), 1106–1121. Retrieved 2017-05-04, from http://www.sciencedirect.com/science/article/pii/S0896627316307772 doi: 10.1016/j.neuron.2016.10.027

Le B, J.-V., Silberberg, G., Wang, Y., & Markram, H. (2007). Morphological, Electrophysiological, and Synaptic Properties of Corticocallosal Pyramidal Cells in the Neonatal Rat Neocortex. Cerebral Cortex, 2204–2213. Retrieved 2012-05-16, from http://cercor.oxfordjournals.org/content/17/9/2204

Markram, H., Muller, E., Ramaswamy, S., Reimann, M., Abdellah, M., Sanchez, C., … Schrmann, F. (2015). Reconstruction and Simulation of Neocortical Microcircuitry. Cell, 163(2), 456–492. Retrieved 2016-11-03, from http://linkinghub.elsevier.com/retrieve/pii/S0092867415011915 doi: 10.1016/j.cell.2015.09.029

Nolte, M., Reimann, M. W., King, J. G., Markram, H., & Muller, E. B. (2019). Cortical reliability amid noise and chaos. Nature Communications, 10(1), 1–15. Retrieved 2019-10-01, from https://www.nature.com/articles/s41467-019-11633-8 doi: 10.1038/s41467-019-11633-8

Perin, R., Berger, T. K., & Markram, H. (2011). A synaptic organizing principle for cortical neuronal groups. Proceedings of the National Academy of Sciences, 108(13), 5419–5424. Retrieved 2012-07-28, from http://www.pnas.org/cgi/doi/10.1073/pnas.1016051108 doi: 10.1073/pnas.1016051108

Ramaswamy, S., Courcol, J.-D., Abdellah, M., Adaszewski, S. R., Antille, N., Arsever, S., … Markram, H. (2015). The neocortical microcircuit collaboration portal: a resource for rat somatosensory cortex. Front. Neural Circuits, 9. Retrieved 2017-04-26, from http://journal.frontiersin.org/article/10.3389/fncir.2015.00044/full doi: 10.3389/fncir.2015.00044

Reimann, M. W., Horlemann, A.-L., Ramaswamy, S., Muller, E. B., & Markram, H. (2017). Morphological Diversity Strongly Constrains Synaptic Connectivity and Plasticity. Cerebral Cortex, 27(9), 4570–4585. Retrieved 2017-09-04, from https://academic.oup.com/cercor/article/27/9/4570/3872356/Morphological-Diversity-Strongly-Constrains doi: 10.1093/cercor/bhx150

Reimann, M. W., King, J. G., Muller, E. B., Ramaswamy, S., & Markram, H. (2015). An algorithm to predict the connectome of neural microcircuits. Frontiers in Computational Neuroscience, 9. Retrieved 2016-11-03, from http://journal.frontiersin.org/Article/10.3389/fncom.2015.00120/abstract doi: 10.3389/fncom.2015.00120

Reimann, M. W., Nolte, M., Scolamiero, M., Turner, K., Perin, R., Chindemi, G., … Markram, H. (2017). Cliques of Neurons Bound into Cavities Provide a Missing Link between Structure and Function. Frontiers in Computational Neuroscience, 11. Retrieved 2017-08-04, from http://journal.frontiersin.org/article/10.3389/fncom.2017.00048/full doi: 10.3389/fncom.2017.00048

Reyes-Puerta, V., Sun, J.-J., Kim, S., Kilb, W., & Luhmann, H. J. (2014). Laminar and Columnar Structure of Sensory-Evoked Multineuronal Spike Sequences in Adult Rat Barrel Cortex In Vivo. Cerebral Cortex, bhu007. Retrieved 2015-07-15, from http://cercor.oxfordjournals.org/content/early/2014/02/10/cercor.bhu007 doi: 10.1093/cercor/bhu007

Ritchie, M., Berthouze, L., House, T., & Kiss, I. Z. (2014). Higher-order structure and epidemic dynamics in clustered networks. Journal of Theoretical Biology, 348, 21–32. Retrieved 2019-05-11, from http://www.sciencedirect.com/science/article/pii/S0022519314000423 doi: 10.1016/j.jtbi.2014.01.025

Rosenbaum, R., Smith, M. A., Kohn, A., Rubin, J. E., & Doiron, B. (2017). The spatial structure of correlated neuronal variability. Nat Neurosci, 20(1), 107–114. Retrieved 2017-03-09, from http://www.nature.com/neuro/journal/v20/n1/full/nn.4433.html doi: 10.1038/nn.4433

Rubinov, M., & Sporns, O. (2010). Complex network measures of brain connectivity: Uses and interpretations. NeuroImage, 52(3), 1059–1069. Retrieved 2019-04-03, from http://www.sciencedirect.com/science/article/pii/S105381190901074X doi: 10.1016/j.neuroimage.2009.10.003

Schreiber, S., Fellous, J. M., Whitmer, D., Tiesinga, P., & Sejnowski, T. J. (2003). A new correlation-based measure of spike timing reliability. Neurocomputing, 5254, 925–931. Retrieved 2015-05-26, from http://www.sciencedirect.com/science/article/pii/S092523120200838X doi: 10.1016/S0925-2312(02)00838-X

Silberberg, G., & Markram, H. (2007). Disynaptic Inhibition between Neocortical Pyramidal Cells Mediated by Martinotti Cells. Neuron, 53(5), 735–746. Retrieved 2013-08-15, from http://linkinghub.elsevier.com/retrieve/pii/S0896627307001110 doi: 10.1016/j.neuron.2007.02.012

Simonyan, K., & Zisserman, A. (2014). Very Deep Convolutional Networks for Large-Scale Image Recognition. 1409.1556 [cs]. Retrieved 2019-04-15, from http://arxiv.org/abs/1409.1556 (1409.1556)

Song, S., Sjstrm, P. J., Reigl, M., Nelson, S., & Chklovskii, D. B. (2005). Highly Nonrandom Features of Synaptic Connectivity in Local Cortical Circuits. PLoS Biology, 3(3), e68. Retrieved 2016-11-03, from http://dx.plos.org/10.1371/journal.pbio.0030068 doi: 10.1371/journal.pbio.0030068

Vogels, T. P., Sprekeler, H., Zenke, F., Clopath, C., & Gerstner, W. (2011). Inhibitory Plasticity Balances Excitation and Inhibition in Sensory Pathways and Memory Networks. Science, 334(6062), 1569–1573. Retrieved 2017-03-29, from http://science.sciencemag.org/content/334/6062/1569 doi: 10.1126/science.1211095

Wang, H.-P., Spencer, D., Fellous, J.-M., & Sejnowski, T. J. (2010). Synchrony of Thalamocortical Inputs Maximizes Cortical Reliability. Science, 328(5974), 106–109. Retrieved 2015-08-04, from http://www.sciencemag.org/content/328/5974/106 doi: 10.1126/science.1183108

Willshaw, D. J., Buneman, O. P., & Longuet-Higgins, H. C. (1969). Non-holographic associative memory. Nature, 222(5197), 960–962.

Yin, W., Brittain, D., Borseth, J., Scott, M. E., Williams, D., Perkins, J., … Costa, N. M. d. (2019). A Petascale Automated Imaging Pipeline for Mapping Neuronal Circuits with High-throughput Transmission Electron Microscopy. bioRxiv, 791889. Retrieved 2019-10-10, from https://www.biorxiv.org/content/10.1101/791889v1 doi: 10.1101/791889

Zhang, D., Zhang, C., & Stepanyants, A. (2018). Robust associative learning is sufficient to explain structural and dynamical properties of local cortical circuits. bioRxiv, 320432. Retrieved 2019-04-26, from https://www.biorxiv.org/content/10.1101/320432v1 doi: 10.1101/320432

